# Reparameterization of the Amber RNA Force Field Non-Bonded Terms

**DOI:** 10.64898/2026.05.18.725894

**Authors:** Anees Mohammed Keedakkatt Puthenpeedikakkal, Chapin E. Cavender, Louis G. Smith, Alan Grossfield, David H. Mathews

## Abstract

All-atom simulations of RNA using molecular dynamics have the promise of modeling conformational preferences, folding thermodynamics, conformational change kinetics, and binding affinities of small molecule therapeutics. These simulations rely on a force field, a set of equations and parameters that model the potential energy as a function of conformation using classical mechanics. One popular force field for RNA is Amber OL3, with the most recent iteration derived in 1999 and with subsequent updates to backbone dihedral parameters. The Amber force field, while frequently used, is known to have limitations; for example, it does not properly stabilize native structures against alternative structures. Here, we provide a new approach to fitting the non-bonded parameters for the force field, specifically atom-centered point charges for electrostatics and the Lennard-Jones parameters. The parameters are fit to quantum mechanics (QM) interaction energies calculated with symmetry-adapted perturbation theory (SAPT), including embedded point charges to represent the electrostatic field from solvent and adjacent nucleotides. In this pilot study with a limited set of fitting data, we use the Amber ff99 equations and atom types unchanged. With the revised parameters, we observe improvement in the stability of native structures relative to alternative structures. Native tetraloop conformations, which unfold with the Amber OL3 force field, are stable on the microsecond timescale with our new force field parameters. We also see improvement in the conformational preferences of tetramers. Crucially, A-form helices are still well-modeled, but we observe additional flexibility in an internal loop that is not consistent with NMR data. Overall, we provide evidence that this new approach to fitting RNA force field parameters to SAPT interaction energies with native-structure context represented as embedded point charges is promising. It offers a flexible solution for revising the equations in future work or for extension to other molecules that interact with RNA, such as proteins and small molecules. We call this new set of force field parameters Amber RNA.ROC26.

## INTRODUCTION

Ribonucleic acid (RNA) can carry genetic information, catalyze reactions, and regulate biological processes^1–4^. It is both an important pharmaceutical target and a therapeutic molecule^5–9^. RNA consists of four nucleobases connected by a sugar and phosphate backbone and can adopt a variety of conformations. Understanding this conformational ensemble is key to characterizing RNA behavior and function.

Molecular dynamics (MD) is a computational tool for studying conformational ensembles, providing atomistic-level resolution and enabling the investigation of equilibrium and ultrafast cellular events^10,11^. This provides an important adjunct to experimental methods^12,13^. An MD simulation trajectory consists of a time series of atomic coordinates propagated from an initial starting structure. Updates to coordinates are defined by a parameterized function, called a force field, that estimates the potential energy of the system as a function of atomic coordinates. The fixed-charge Amber^14^ force field consists of terms for bonds, angles, dihedrals, and nonbonded interactions:

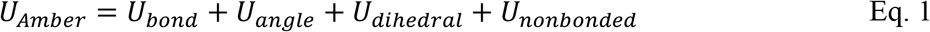

Modern Amber force fields inherit many parameters from the second generation force field parm94^15^, which provided a framework for MD simulations of RNA. Parm99 improved the simulation stability by refining the χ dihedral angles to match *ab initio* calculations and recalibrated so that sugar puckers maintain the canonical A-form geometry. Perez et al.^16^ reparametrized α and γ dihedrals to allow stable microsecond scale simulations of DNA with more complete model systems. Zgarbova et al. modified the glycosidic torsions χ of purines and pyrimidines to reduce ladder-like distortions in RNA MD simulations^17^.

In the current force field, bond and angle parameters are modeled by harmonic potentials:

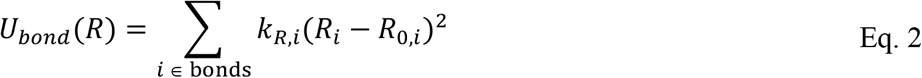

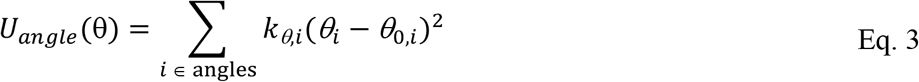

where the energy constants, *k*_*R,i*_ and 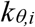, were determined by fitting to experimental results of model compounds using normal mode analysis. Equilibrium geometries (*R*_*0*_ and *θ*_*0*_) were derived from X-ray structures and microwave spectroscopy^18^. Bond and angle stretch force constants (*k*_*R,i*_ and 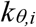) were derived from infrared and microwave spectroscopy^18–20^.

The nonbonded parameters of the RNA Amber force field include terms for electrostatics (modeled as atom-centered point charges) and van der Waals terms:

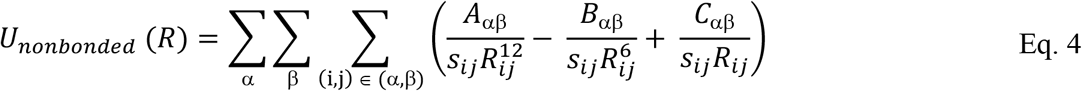

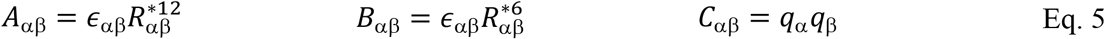

where *R*_*ij*_ is the distance from atom *i* to atom *j, ϵ*_αβ_ is the LJ well depth between atom type indices α and β, R^*^_αβ_ is the half radius, *q* is the partial charge on atom centers and *s*_*ij*_ is the 1-4 scaling factor for nonbonded interactions. These values are largely unchanged since ff99^21^.

The van der Waals interactions are approximated with a Lennard-Jones (LJ) potential (terms 1 and 2 in eq. 4). These parameters were taken from the optimized potentials for liquid simulations (OPLS) approach, in which parameters were manually fit such that densities and heats of vaporization from short simulations of neat liquids matched experiments^15,22^. The neat liquids used here were small molecule analogs of biomolecules, such as N-methyl acetamide, tetrahydrofuran, and pyridine. It was assumed that these results can be transferred to RNA atoms in an aqueous environment.

Atom centered point charges were fit by the restrained electrostatic potential (RESP) method^23^. Residues were placed in a grid and the electrostatic potential (ESP) on the grid was estimated using Hartree-Fock with a 6-31G* basis set. The point charges were then fit to reproduce the gas-phase ESP values on the grid using RESP, which restrains the magnitudes of the charges on buried atoms to small magnitudes^23^. To compensate for the lack of solvent, HF/6-31G* was used, which was known to overpolarize wavefunctions to mimic an aqueous environment^21^. RESP fitting has been noted to be difficult to reproduce, which in part arises from differences in fits when a residue is oriented differently in the grid^24^ and from differences in intramolecular conformations^25^.

Torsional (dihedral) energies are represented as a sum of cosine terms:

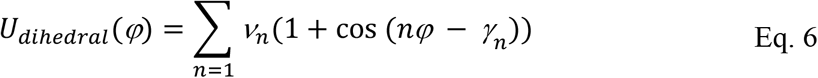

where *φ* is the dihedral angle, *n* is the multiplicity, *γ*_*n*_ is a phase offset and *ν*_*n*_ is the barrier height. Dihedral parameters were fit using small organic models to either QM energies using MP2/6-31G* or to experimental data, when available^15^. These terms were fit last in the force field, as they have no single physical potential to which they correspond.

As computational resources improved and microsecond-scale simulations became possible (and eventually routine), the shortcomings of these force fields became apparent^10,11^. While helices are generally stable, native structures of hairpin and internal loops disappear in simulations on the μs timescale^26–30^. Extensive simulations of RNA tetramers found that simulations largely sample conformations that are not supported by NMR experiments^31–37^.

Multiple efforts attempted to resolve these issues by improving force field parameters. Most modifications were focused on subsets of dihedral parameters, using fits to QM calculations, and have resulted in better agreement with NMR data^17,29,38–40^. There have been attempts to improve nonbonded parameters^41–44^, but transferability has proven to be limited, or these force fields did not resolve the inability of previous force fields to stabilize native conformations against alternative conformations. Recently, remarkable progress was demonstrated for UUCG tetraloop simulations, where three improved Amber force fields^35,37,43^ were shown to preserve the native structure for multiple 20 μs simulations.^36^ Folding free energy change calculations in that study, however, showed insufficient stability against alternative conformations^45^.

In this work, we present a new approach to fitting nonbonded parameters for a fixed-charge force field. We focused on the non-bonded parameters because the bonded parameters are more closely rooted in experimental data. The atom-centered partial changes and Lennard-Jones well depth and half-radius parameters are fit to average QM interaction energies for condensed-phase and vacuum systems (Figure 1). A diverse dataset of RNA conformations is curated according to sequence and conformation, including base pairing, base stacking, base-phosphate, and base-sugar interactions. The QM interaction potential energies of these structures are computed in the presence of point charges representing the solvent environment and neighboring nucleic acid atoms, inspired by the IPolQ method^46^. The charges are fit, followed by a fit of the LJ well depth and half-radius parameters. The backbone and glycosidic dihedral parameters are then refit using the method of Aytenfisu et al^29^. A second iteration is performed to improve the self-consistency of the new charges and LJ parameters to solvent configurations. In this study, we focus on the current Amber ff99 equations and the current set of adjustable parameters. The approach, however, is flexible and could be used to fit parameters for other functional forms or to extend the parameter set, such as adding additional atom types for LJ parameters (Figure S1).

The newly fit force field parameters, which we call RNA.ROC26 (for Rochester), were tested using MD trajectories of tetraloops, duplexes, and tetramers, i.e. four nucleotide long RNA. These simulations showed improved agreement with experimental structures on the μs timescale, with the one exception of a specific internal loop motif.

## RESULTS

### Curating a diverse data set of training structures

The first step in the fitting procedure (Figure 1) was assembling a diverse set of training conformations as illustrated in Figure 2. To have good quality parameter fits, the training conformations need to represent a diversity of sequences and conformations. Our procedure is designed to accomplish this by identifying a large candidate set and then clustering from this set to identify a variety of representative conformations.

To start, we downloaded the coordinates for all structures in the Representative Set of RNA 3D Structures database from Bowling Green State University^47^. This set consists of non-redundant RNA structures from the Protein Data Bank^48^ (PDB). All interacting RNA nucleotide pairs were extracted from the 1371 nonredundant RNA structures, where interacting residues were defined as being either consecutive in sequence or engaged in stacking or base pairing interactions. The distance between the O3’ of the 5’ residue and the P atom of the 3’ residue was used to identify consecutive nucleotides. The cutoff was set to 2.32 Å based on the distribution of O3’-P distances. Pairing or stacking nucleobases were identified using the anisotropic position vector used by Bottaro et al.^49^. All RNA pairs with the magnitude of the anisotropic position vector less than √2.5, the same threshold used by Bottaro et al., were considered interacting and selected for clustering. All interacting RNA pairs in each structure in this set were considered as candidate conformations for the training dataset.

We then used clustering to identify a diverse dataset of interacting nucleotide pairs. We developed a distance metric based on geometric contributions to nonbonded interactions between pairs of heavy-atom types, referred to as the pairwise distance matrix (Methods). Briefly, the pairwise distance matrix includes terms of *R*^*-1*^, *R*^*-6*^, and *R*^*-12*^, where R is the heavy-atom-heavy-atom distances between nucleotides. This pairwise distance matrix weights the atomic distances according to their usage in the force field’s non-bonded terms (Eq. 4). Using a Euclidean distance based on the columns of this pairwise distance matrix, density peak clustering^50^ was used to cluster the candidate nucleotide pairs (Methods).

The atom types present in each nucleotide pair conformation depend on the interacting nucleobases, so the set of candidate conformations was partitioned into ten groups based on nucleobase identity (AA, AC, AG, AU, CC, CG, CU, GG, GU, and UU) and clustered separately within each group. From each group, 32 cluster centers were selected to form the training set. The results of density-peak clustering are provided in the SI, along with the PDB IDs of selected structures (Table S1).

To include additional diversity in the training set, candidate conformations with nucleobase-phosphate and nucleobase-sugar interactions were extracted from the *Haloarcula marismortui* large subunit ribosomal RNA (PDB ID 1S72, Chain 0) and added to the training set (Methods)^51,52^.

### Simulation of Required Sample Size

We tested our ability to identify diverse training conformations by fitting Amber ff99 non-bonded parameters to Amber ff99-determined interaction energies^21^ to training sets of different sizes. To understand the dependence of the fit quality on the size of training dataset, the number of conformations selected from each of the ten sequence groups was varied from 3 to 100. For each size, a clustered training dataset was constructed by density peak clustering. Additionally, ten random training datasets were constructed by randomly sampling candidate conformations without replacement. These random datasets represent negative controls. The vacuum Amber ff99 nonbonded interaction energy was used as the parameter fitting target. We calculated the root mean squared error for the interaction energies estimated by the fit parameters for a testing set consisting of all candidate conformations not included in the corresponding training set.

This test demonstrated that the clustering was able to sample a diverse and representative set of conformations (Figure S2). We found that a training set with 130 conformations was able to reproduce ff99 nonbonded parameters, slightly more than the 116 parameters to fit. When the number of conformations is lower than the number of parameters, the errors in the estimated interaction energies are large because the fit is underdetermined. The fitting dataset derived by clustering rapidly works well for fitting parameters as a function of training conformation size once the number of conformations exceeds the number of parameters.

When using the QM interaction energies as the fitting target, we expected more conformations would be needed because the Amber ff99 functional form only approximates the QM landscape. To balance the need for an adequate number of fitting targets and the computational cost, a total of 384 conformations were used. The set includes 32 each from groups based on nucleobase identities, 32 base-phosphate and 32 base-sugar interactions.

### MD simulation to sample solvent electrostatic field

To represent the effect of solvent and context accurately in our training data, we chose an approach inspired by the IPolQ method^46^ where we immersed the model systems in atomistic water and salt, generating simulated mean densities of solvent charge using Molecular Dynamics (MD; Figure 3). Each training conformation includes the primary residue and additional residues that represent the local RNA structure. To include local RNA structure, residues that interact with either of the selected primary residues are also included (Figures 3A and S3A). These interacting residues are determined using the same criteria mentioned above. Each of the 384 structures in the training set were solvated in a truncated octahedron of OPC water^53^ with 40 Å of margin. K^+^ counterions were added to neutralize the total charge, and K^+^/Cl^-^ ions were added using the SLTCAP method to achieve a bulk salt concentration of 150 mM^54,55^.

To determine how much solvent was needed to converge the mean solvent electrostatic field, we ran pilot simulations on a validation set of conformations constructed from the exemplars of the four most populated clusters for each sequence pair (Table S2). A 40 ns MD simulation with a 40 Å solvent margin was used as a reference to calculate the root-mean-square error (RMSE) of electrostatic energy (ESE) across all validation set conformations. We chose the 40 Å margin to attempt to include solvent past which counterion densities are not perturbed by the RNA^56^. The root-mean-square error (RMSE) for the solvent electrostatic potential energy on all the RNA atoms as a function of simulation length and solvent cutoff box size (Figure S4) was used to determine the adequate box size and simulation length for sampling the solvent environment. The RMSE across all RNA atoms in the validation set is 5.8 kcal/mol for the first 2 ns, and the RMSE decreased to 1.6 kcal/mol by 20 ns. Similarly, the RMSE in ESE as a function of cutoff for solvent was 52 kcal/mol at 5 Å, relative to the full 40 Å simulation size. A similar trend was observed for 20 ns simulations. Therefore, a 20 ns simulation length and 40 Å margin of solvent were chosen. For these choices of simulation length and solvent margin, the RMSE introduced here were less than 0.1% of the total ESE, which are on the order of -4×10^5^ kcal/mol.

### Image charge representation of the solvent electrostatic field

A mesh of image charges was used to approximate the mean electrostatic field due to solvent atoms far from the RNA, following the implicitly polarized charges (IPolQ) method used to fit charges for amino acids^46^. In this method of images, a complicated distribution of charges producing an electrostatic potential (ESP) throughout a volume of space was represented as a distribution of point charges on the bounding surface of that volume, called image charges, that can produce the same ESP (Figure 4; Methods).

### Point Charges to represent local RNA structure

To model the influence of local RNA structure on the electrostatic environment, we reasoned that we needed to model the polarization induced by RNA nucleotides adjacent to the interacting pair. Thus, we included these adjacent nucleotides in the QM system as point charges to model the influence of local RNA structure. For nucleotides pairs not covalently bound to each other, the QM system comprised all atoms from the nucleotides except phosphorus and non-bridging oxygens. To build a closed shell system for SAPT calculations, i.e., each occupied molecular orbital contains exactly two paired electrons with opposite spin, the covalent bonds O5’ and O3’ to phosphorus were removed, and hydrogens replaced the phosphorus (Figure S4). This ‘capping’ strategy was used throughout to avoid including phosphate atoms in the QM systems, because it has many electrons and is therefore expensive to represent accurately. The remaining atoms for adjacent nucleotides were represented as point charges (Figures 3B and S3B). The Amber ff99 phosphorus atom charge was distributed evenly to the charges representing the three non-QM atoms covalently bonded to the phosphorus (nonbridging phosphate oxygens and either O5’ or O3’).

For conformations in the training dataset with covalently bound interacting nucleotides, the nonbridging phosphate oxygens and P were removed from the 5’ end as was done for non-adjacent residues, and O3’ was truncated as OH. As the 5’ nucleotide and 3’ nucleotide are covalently linked by phosphodiester bonds, an extension of SAPT0 called intramolecular SAPT (I-SAPT)^57^ was used to compute the QM interaction energy.

For SAPT calculations of training conformations with a base-phosphate interaction, the nucleotides covalently bound to the phosphodiester group were included in the QM system (Figure S5D). The 5’ end of the 5’ nucleotide and the 3’ end of the 3’ nucleotide were truncated with hydrogen, and other adjacent RNA nucleotides were included as point charges to represent local RNA structure. For the nucleobase interacting with the phosphate, the above-mentioned non-covalently bound nucleotide representation (Figure S5B) was used for QM calculations. The nonbonded parameters for the nonterminal C5’, C3’, and phosphate atoms were informed only from the nucleobase-phosphate interactions in the training conformations.

The above-mentioned representation of nucleobase-nucleobase interaction was extended to depict the nucleobase-sugar interactions, depending on whether two residues containing the nucleobase and sugar are covalently bound (Figure S5D) or not (Figure S5B).

Figures 3C and S3C illustrate the final point-charge representation of an RNA conformation in the training set for a non-covalently bound and a covalently bound interacting nucleotide pair, respectively.

### QM interaction energy calculations

We chose to use symmetry-adapted perturbation theory (SAPT) to determine interaction potential energies, therefore the non-bonded parameter fits were for intermolecular interactions, and bonded terms did not enter the fitting calculations^58^. For each training structure, the point-charge representation (Figures 3C and S3C) was used to calculate QM interaction energies using electrostatically embedded symmetry-adapted perturbation theory (EE-SAPT)^59^, truncated to the lowest order with modified exchange scaling (sSAPT0), using the jun-cc-pVDZ basis set^57,60^. This model chemistry reproduced the gold-standard interaction energies computed with coupled-cluster perturbative triples (CCSD(T)) in the complete basis set with a mean absolute error of 0.49 kcal/mol^57^. The accuracy of sSAPT0/jun-cc-pVDZ is comparable to second-order Moller-Plesset perturbation theory MP2/aug-cc-pVTZ and density functional theory (DFT) methods like B97D3/aug-cc-pVDZ^57^. sSAPT0 provided inter-residue interaction potential energies that correspond to only the nonbonded terms in the force field.

The convergence of the QM interaction energies as a function of image mesh radius was evaluated by comparing the validation set (See “MD simulation to sample solvent electrostatic field” above) potential energies obtained with the mesh to those obtained with all solvent atoms as explicit point charges (Figure S6). Compared to QM interaction energies that included explicit point charges for all atoms in solvent molecules within 30 Å of any RNA atom, the QM interaction energies computed using image charges on a mesh with a 4 Å radius, replacing solvent molecules located further than 4 Å but within 30 Å of any RNA atom, the RMSE was less than 1×10^−3^ kcal/mol. Based on this comparison, the image mesh radius of 4 Å was used to approximate the mean electrostatic field.

For each structure in the training set, two sets of QM potential energies were calculated: One without the solvent point charges and one in the presence of the solvent point charges, i.e., a condensed-phase representation, as explained above. The local RNA environment is included as point charges in both cases. For fitting, the average interaction energies of these two calculations were used. We chose this specific approach, one that averages the potential energies before fitting, because we are fitting both LJ parameters and charges in a single procedure with a goal of having self-consistent parameters with the best approximation of interaction energies.

### Fitting nonbonded parameters to QM interaction energies

Given the conformations and their interaction potential energies, our goal was to fit the nonbonded force field parameters to best reproduce interaction energies determined by QM (Figure 1). In the Amber force field model (eq. 1), the nonbonded energy terms exhibit a nonlinear dependence. The nonbonded fits were done in two steps, where each step used a Monte Carlo search with a local nonlinear optimizer. Atom-centered point charges were fit first, followed by the Lennard-Jones (LJ) parameters, i.e. half-radius and well depth. In each fit, L2 regularization was employed to reduce the risk of overfitting parameters for this relatively small set of training conformations.

The cost function for fitting partial charges to the fitting target was:

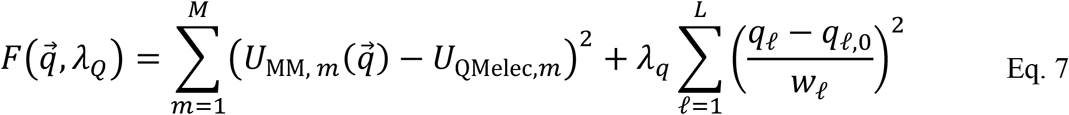

where M denotes the total number of training conformations and L represents the number of charge atom types. Each atom is treated as a distinct atom type for partial charges, yielding a total of 76 charge types in RNA (following Amber ff99^21^). 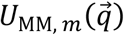 is the electrostatic component of interaction energy calculated using eq. 4 and *U*_QMelec,*m*_ is the electrostatic component from the sSAPT0 interaction energy for training conformation *m*. The first term represented the sum of squared residuals (SSR) between 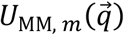 and *U*_QMelec,*m*_, where 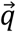 was the fit. The electrostatic components from the average QM energies were used as the fitting target (*U*_QMelec,*m*_). To obtain *U*_QMelec,*m*_, the LJ component of the force field was calculated using the Amber ff99 nonbonded parameters^21^ and subtracted from the average QM energy.

The next term in the cost function was the regularization term; *λ*_*Q*_ determines the strength of regularization for partial charges. Each regularization term had a regularization target, *q*_*ℓ*,0_, representing the mean of the Gaussian priors and a weight representing the standard deviation of the prior, *w*_*l*_. For fitting the partial charge, the Amber ff99 charges were used as the regularization target, and 1 e^-^ was used as *w*_*ℓ*_. This regularization term applied harmonic restraints in the fit that are analogous to RESP, which uses hyperbolic restraints that are restrained towards zero^23^.

The cost function for fitting LJ parameters to the fitting target was:

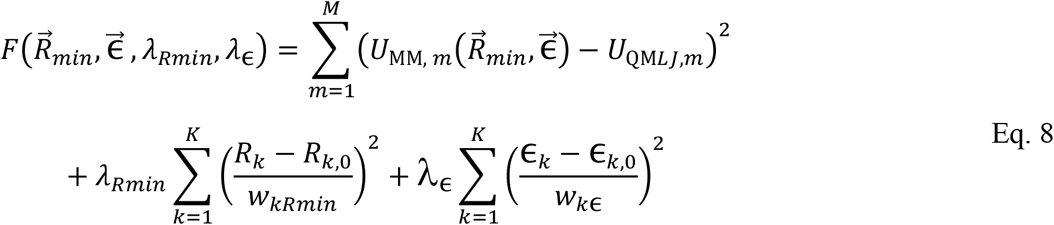

where K denotes the total number of LJ atom types. Amber ff99 has 14 LJ atom types^15^, and the same representation was used for the LJ parameter fit. 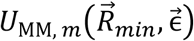 is the LJ component of eq. 4 and *U*_QM*LJ,m*_ is the LJ component sSAPT0 interaction energy. In the LJ fit cost function, the first term represents the SSR between the LJ energy term, 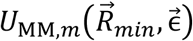, and the target fitting energy, *U*_QM*LJ,m*_. Here, *U*_QM*LJ,m*_ was calculated by subtracting the electrostatic energy term 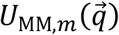 from the average QM energy. The electrostatic energy is calculated using the newly fit partial charges. The second term is a regularization term where *λ*_*Rmin*_ and *λ*_ϵ_ are regularization strengths. The regularization targets (*R*_*k*,0_, ϵ_*k*,0_) were the mean of the nonzero Amber ff99 parameters grouped by element, except for two types of hydrogen, polar and nonpolar (Table S3), which were separated into two “elements” for the purpose of calculating means. The regularization weight (*w*_*k*R_, *w*_*k*ϵ_) for the VDW half radii was 1 Å and 0.1 kcal/mol for the VDW well depth.

The regularization strengths (λ) were determined by 8-fold cross-validations using a grid search to identify values that minimize the SSR (the first terms in equations 7 and 8), excluding the regularization terms. This ensures that regularization strengths are chosen that allow improved agreement with the QM potential energies without overfitting to the dataset. For each λ in the cross validation, the cost function was minimized via gradient decent. This is efficient and precise, but it converges to the nearest local minimum to the starting point, making it sensitive to initial starting estimates. To address this, multiple starting estimates were used (Methods).

Cross-validation of the charge λ_q_ had minima at λ_q_ = 10^2.500^ and λ_q_ = 10^2.125^ for the first fitting iteration and second iteration, respectively (Figure 5A, 5C). The SSR from LJ cross-validation shows a broad, flat minimum (Figure 5B, 5D). A low constant (λ_Rmin_, λ_ϵ_ < 10^0^) minimized the SSR, but risked overfitting (Figure S7). λ_Rmin_=10^1.250^ and λ_ϵ_ = 10^0.250^ were selected for the LJ parameter fits in iteration 1 and λ_Rmin_=10^1.250^ and λ_ϵ_ = 10^1.875^ for iteration 2. These λ values lie in a tradeoff region with near-optimal SSR to the unrestrained fits (i.e., with no regularization). We chose the nonzero log(λ) values to ensure that parameters are not overfit to the dataset.

Once the regularization strengths were determined, the basin-hopping algorithm was used to search for the global minimum of each cost function (eq. 7 and then eq. 8). Basin hopping is a global optimization algorithm. A random vector perturbs the parameters, followed by a local gradient-descent optimization is performed, and the new minimum is accepted or rejected based on the Metropolis-Hastings criterion, using the cost function values at the new and previous minimum. This continues for a fixed number of steps, and then the parameter set with lowest cost function in the sampled sets is taken as the solution. This basin hopping approach allows the algorithm to escape local minima and explore the broader parameter space in search of a global minimum. With sufficiently large perturbations, the optimization can cross large barriers in the fit landscape. A Gaussian random variable with zero mean and standard deviation equal to one-tenth of the range of parameter boundaries was used to perturb the parameters. The vector was multiplied by a step size parameter that was adaptively adjusted based on the acceptance rate during the optimizations.

### Iterating the fitting pipeline to improve nonbonded parameters

Because the charges and LJ parameters were fit separately, we chose to iterate to improve the quality of the fit. The first set of fit charges and LJ parameters were used to run the solvent MD simulations to resample the solvent environment. Then, the QM potential energies were recalculated, and the fit was repeated using cross-validation and basin-hopping steps (Figure 1).

The fit parameters (Figure 6) from both iterations are given in Table S4. During basin-hopping with the full fitting conformation dataset, the SSR without the regularization terms for the charges improved from 3923 kcal^2^/mol^2^ for the Amber ff99 parameters to 916 kcal^2^/mol^2^ and 619 kcal^2^/mol^2^ for the first and second iteration, respectively. Likewise, the LJ parameters SSR improved in the fit from 801 kcal^2^/mol^2^ to 581 kcal^2^/mol^2^ and then to 548 kcal^2^/mol^2^.

### New dihedral terms for fit nonbonded parameters

After the nonbonded parameters were fit, the dihedral terms must be refit because they represent a correction to the other terms to ensure a final torsional preference. We used the method of Aytenfisu et al.^29^ to refit backbone dihedral parameters after each iteration. In this approach, multiple linear regression was used to fit the parameters to potential energies calculated with the dispersion-corrected density functional B97, the damping function D3(BJ), and the AUG-CC-PVTZ basis set^61,62^. The amplitudes and phases of the backbone dihedral angles α, β, γ, ϵ, ζ and the four glycosidic torsions χ were simultaneously refit. A dataset of ∼31 thousand structures, which includes structures from the PDB and dihedral scans of nucleoside conformations, taken from Aytenfisu et al.^29^, was used for the fit. The dihedral potential energy profiles using the new fit dihedral terms are in Figure S8.

### Explicit solvent simulations to test the fit parameters

To test the new nonbonded parameters, 2.5 μs simulations were run in quadruplicate with RNA tetraloops and duplexes (Table 1). These structures represent a variety of well-studied systems. We expect that the A-form helices, 2JXQ and GC duplex, will remain close to their starting structures for all force fields. The AU duplex is a negative control because it has melting temperature similar to the temperature at which the simulations are run. Finally, the internal loop, 2DD2, has a structure determined by NMR and has been studied by us previously by MD.

**Table 1:**
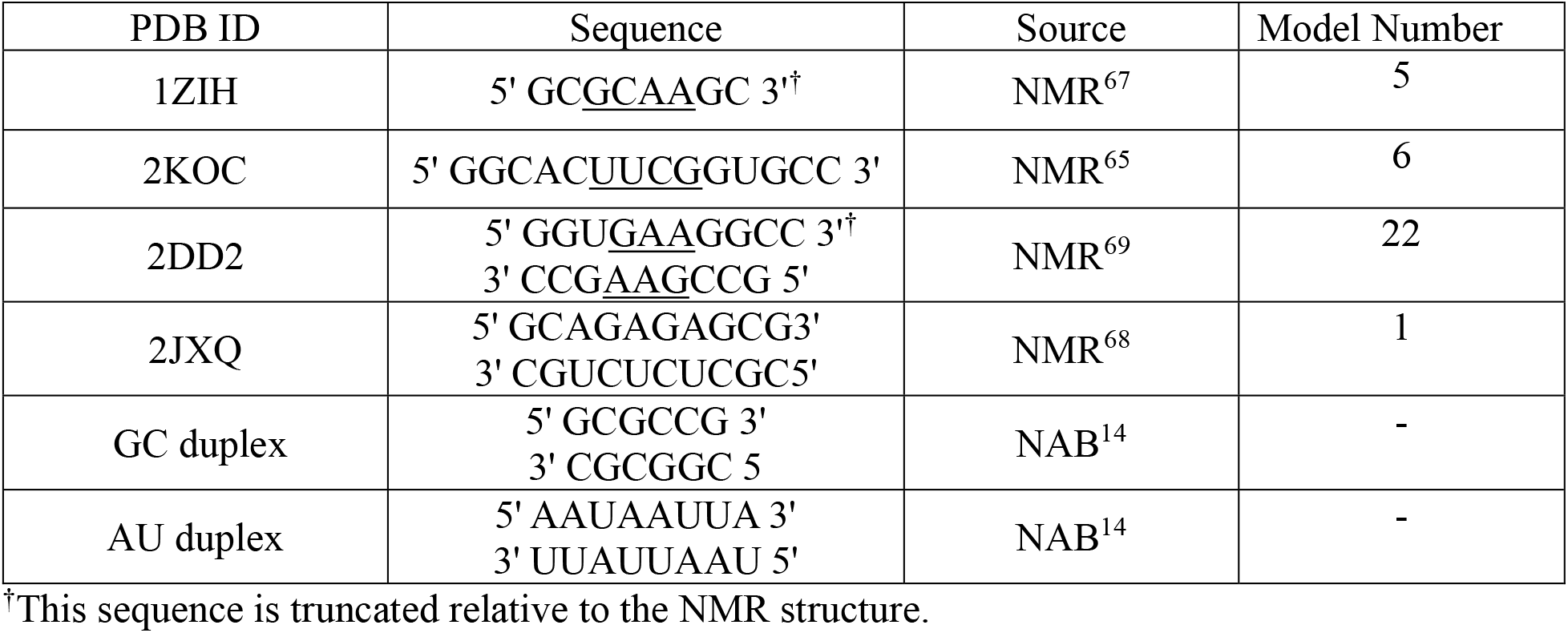
Structures used to run benchmark MD simulation. Underlined nucleotides are in hairpin or internal loops. NAB for source means the structure were constructed in A-form geometry using NAB^14^. For NMR structures, we used the model with lowest potential energies after energy minimization^29^.

The heavy atom RMSD relative to their starting solution structures are given in Figure 7 (hairpin loops) and Figure 8 (duplexes). For comparison, simulations were also run with the Amber ff99 + bsc0 + χ_OL3_ force field^16,17,63^ and RNA.ROC^29^ force fields (Figure S9).

The 1ZIH and 2KOC structures contain the GCAA and UUCG tetraloops, respectively, which serve as standard benchmark systems for RNA force fields. The tetraloops are common motifs in tertiary structures of biological RNAs^64^. The loop residues adopt well-defined structures stabilized by both base-phosphate and base-sugar interactions that are poorly described by the OL3 forcefield^17,65,66^.

For the tetraloop benchmark simulations, RNA.ROC26 produced relatively stable simulations, while the Amber ff99 + bsc0 + χ_OL3_ force field^16,17,63^ generally did not maintain the solution NMR structure. In previous studies^22^, the UUCG tetraloop of 2KOC^65^ is not stable for μs scale simulations, but with RNA.ROC26, the tetraloop is stable up to 2.5 μs (Figure 7B); Amber ff99 + bsc0 + χ_OL3_ denatured the NMR structure and RNA.ROC had substantial fluctuations after 1.5 µs. The GCAA tetraloop^67^ also stayed close to the experimental structure with RNA.ROC26 relative to the other two force fields. One simulation, however, had a substantial increase in RMSD after 1.5 µs because the tetraloop’s A5 bulged out of the hairpin during the simulation, which is not found in NMR experiments^67^. At about 2.5 µs, A5 returned to the initial NMR configuration.

To run simulations of Watson-Crick-Franklin helices, the 2JXQ^68^ system and GC duplex systems were used, which A-form structures. The GC duplex structure was generated using NAB^14^. 2JXQ and GC_duplex simulations (Table 1) showed comparable results across the three force fields (Figure 8 and S10), with heavy atom RMSD less than 2.5 Å across the simulations.

The 2DD2 system contains a 3×3 internal loop (GGA/AAG) with a central trans Hoogsteen/Sugar Edge A-A pair enclosed between two trans Hoogsteen/Sugar Edge A-G pairs. NMR demonstrated that the central A-A pair exchanges between two alternative pairs on the tens of μs timescale that differ as to which of the two adenines provide the Hoogsteen or sugar edge^69^. We previously studied the pathway for this exchange by MD simulation using nudged elastic band and umbrella sampling^70–74^.

In the internal loop benchmarks, the pairing of the central A-A pair was unstable with RNA.ROC26, with one of the adenines bulging out into the solvent during simulations and then returning to the pair (Figure 8 and S10). However, this pair was stable in the Amber ff99 + bsc0 + χ_OL3_ force field and RNA.ROC simulations (Figure S9 and S10). For all three force fields, the G-A pairs at either end of the internal loop were stable. For this system, the heavy-atom RMSD for internal loops, including the closing base pairs, showed larger fluctuations for RNA.ROC26 than in Amber ff99 + bsc0 + χ_OL3_ force field and RNA.ROC (Figure S10). The large RMSD for RNA.ROC26 was inconsistent with the NMR structure.

The AU duplex, composed of only AU pairs, was designed as a negative control. The melting temperature when using a salt correction is estimated as 302 K^75,76^, and therefore the duplexes should show substantial fluctuation or denaturation for simulations at 300 K. It was completely denatured in 3 of 4 RNA.ROC26 simulations by 0.25 µs, and similar results were observed for the other two force fields later in the simulations.

### Simulations of tetramers

RNA tetramers are a good system for benchmarking nonbonded interactions and are well-studied by NMR and by prior simulations^32^. Their small size makes sampling easier and faster. The NMR structures of AAAA^31^, CAAU^31^, CCCC^38^, GACC^33^, and UUUU^31^ are close to A-form-helix conformations with some deviations. For example, NMR NOE experiments of AAAA, CAAU and GACC show that, at least some of the time, the sugar of the 3’ nucleotide is inverted. MD simulations using the Amber ff99 + bsc0 + χ_OL3_ force field^16,17,63^ yielded intercalated structures, where the base in position 1 inserts between bases 3 and 4, for which there is no support in the NMR NOEs. These aberrant structures often emerged within the first microsecond of a simulation; once found, they often remained for the duration of the simulations. Except for GACC, Amber ff99 + bsc0 + χ_OL3_ simulations, a majority of the structures observed were intercalated^31,32^.

In this work, we ran explicit solvent MD simulations of AAAA, CAAU, CCCC, GACC, and UUUU using RNA.ROC26, Amber ff99 + bsc0 + χ_OL3_, and RNA.ROC force field. For each sequence, five independent simulations were run for 2 μs each. Following Bergonzo & Cheatham^32^, we focused our analysis on RMSD to the A-form structure and clustering the conformations. Figure S11 displays the histograms of RMSD for each simulation and Figure 9 displays the histograms aggregated across the simulations.

In general, RNA.ROC26 showed a larger population of low RMSD structures than the other force fields. A-form like conformations were well populated when using RNA.ROC26. However, for AAAA, the A-form like population was low and ∼30% of the structures showed intercalation (Figure 9A:F3). Similar to NMR observations, the RNA.ROC26 simulations showed a population of inverted sugars for the 3’ nucleotide (Figure 9A:F2, 8B:F2; 8D:F2). For AAAA, the RNA.ROC force field simulations were similar to RNA.ROC26. For CAAU^31^, RNA.ROC26 favored an A-form-like orientation, whereas RNA.ROC favored the structure with an inverted 3’ sugar (Figure 9B:R1) and Amber ff99 + bsc0 + χ_OL3_ yielded a majority of intercalated structures (Figure 9B:O1). In GACC^33^ simulations, all three force fields populated both A-form and inverted sugars (Figure 9D). CCCC^38^ simulations demonstrated two major clusters, one A-form like and another with the inverted 3’ terminal sugar (Figure 9C). The RNA.ROC26 forcefield favors the A-form-like structure and ROC favors the structure with an inverted sugar, but Amber ff99 + bsc0 + χ_OL3_ favors intercalated structures. UUUU simulations structures were mostly A-form like for RNA.ROC26 simulations (Figure 9E: F1), where Amber ff99 + bsc0 + χ_OL3_ formed mostly intercalated structures (Figure 9E: O3) and RNA.ROC preferred the inverted 3’ terminal sugar (Figure 9E: R1).

## DISCUSSION

In this work, we present a new parameterization of the Amber force field, RNA.ROC26. The partial charges, LJ well depth, and half radii are fit to QM energies of a diverse set of training conformations. In contrast, the standard LJ parameters used in Amber ff99^21^ were inherited from OPLS^22,77^. Benchmarking studies have consistently shown that the Amber force field overestimates the strength of stacking interactions^27,32,66,78^. The neat liquid simulations in the OPLS model were performed using small-molecule analogs of functional groups in biomolecules^15,18^. However, the aqueous environment around RNA differs significantly from that in those solvents, especially due to the presence of the negatively charged phosphodiester group and the 2’-OH group. The stacking and hydrogen bonding interactions within the bases themselves should also be sensitive to their average polarization, as they are aromatic moieties. We hypothesize this is one of the reasons for the simulation artifacts with the Amber ff99 + bsc0 + χ_OL3_ force field. In our approach, converged MD simulations of training conformations in OPC water were used to represent a condensed-phase environment.

Chen and Garcia folded the UUCG, GCAA, and CUUG tetraloops by uniformly reducing the LJ σ of base-base interactions to match close contact distances observed from high-level dispersion-corrected coupled cluster theory calculation of gas-phase nucleobases^41^, and the well depth ϵ was adjusted to achieve the correct stacking propensity in solutions. The adjusted σ values for carbon and nitrogen match the half-radii observed in this work, but the RNA.ROC26 oxygen half-radius is marginally larger by ∼0.2 Å. The RNA.ROC26 LJ well depth are found to be larger in magnitude than that of Chen and Garcia^41^ by ∼0.03 kcal/mol for carbon, nitrogen, and oxygen.

In another approach^42^, a subset of nonbonded parameters was revised to improve the description of base pairing and base stacking. The fit charges in RNA.ROC26 are slightly larger in magnitude for adenine and cytosine than the parameters from Tan et al.^42^, whereas for uracil and guanine, they are lower in magnitude. N4 of cytosine and N6 of adenine shows highest deviations of ∼0.1 e-. The RNA.ROC26 half radii are consistent with Tan et al. ^42^, except for hydrogen. The LJ half-radius and well depth of hydrogen were set to zero in Tan et al. ^42^. In contrast, RNA.ROC26 hydrogen well depth minima were kept at the ff99 value, 0.0157 kcal/mol in the fitting procedure.

Steinbrecher et al^79^, address the imbalance in the electrostatic interactions between water and phosphate oxygens by increasing the phosphate oxygen’s LJ half radius relative to Amber ff99^21^. The resulting radii are higher than RNA.ROC26 by 0.05 Å for deprotonated phosphate oxygens (O) and 0.16 Å for protonated oxygens (OH). In contrast, the bridging oxygen (OS) radius are lower than RNA.ROC26 by 0.06 Å.

Fröhlking et al.^43^ proposed an approach to ‘patch’ the issues with Amber forcefields by introducing a generalized Hydrogen-Bond fix (gHBfix), which is an additional energy term applied to specific atom pairs involved in hydrogen bonding. Instead of manually guessing which bonds to strengthen or weaken, they used an automatic optimization framework to tune parameters to match experimental data, specifically by minimizing the discrepancies to solution NMR observables like ^*3*^*J* scalar coupling, NOE intensities, and the experimental native state populations of RNA tetramers and tetraloops.

Raguette et al.^35^ found that CH-O interactions are overly repulsive in the Amber force fields by comparing data extracted from high-resolution RNA crystal structures to energy profiles from QM and force field calculations. They implemented a simple and targeted adjustment of CH-O repulsive interaction by increasing the *R*_*min*_ of oxygen atoms relative to Amber FF99^21^. The *R*_*min*_ values in RNA.ROC26 are lower than Raguette et al.^35^ by 0.05 Å for O and 0.08 Å for OH. The OS half radius is larger for RNA.ROC26 by 0.6 Å. The oxygen half radii reported by Raguette et al.^35^ and Steinbrecher et al^79^ show trends similar to RNA.ROC26 half radii relative to Amber ff99.

The tetramer simulation benchmarks with RNA.ROC26 showed a low population intercalated structures, except for AAAA, where the population is ∼30%. Compared to OL3^21^, this is a significant improvement, but still not perfect agreement with the NMR data. RNA.ROC26 populated both A-form-like structures and other NMR-observed structured conformations. The UUCG and GCAA tetraloop simulations were stable with the ROC.ROC26 force field, where OL3 demonstrated denaturation. In the case of 2DD2, the new parameters were unable to stabilize the internal loop, but the RNA.OL3^17^ and RNA.ROC^29^ force field simulations were stable, indicating scope for improvement in the RNA.ROC26 parameters.

In RNA structures, nucleotides predominantly interact through base pairing or base stacking. The influence of these interactions is absent during the RESP charge fitting^23^. The partial charges derived from RESP fit reproduced the electrostatic potential on grid points around nucleosides in the gas phase. To compensate for the lack of solvent, RESP uses the Hartree-Fock (HF/6-31G*) method^80,81^, which systematically overpolarizes the electron density to mimic the electrostatics of a condensed phase^15^. This approach might reproduce general trends in the electrostatic field in the condensed phase. However, the polarization induced by charged species, such as the phosphodiester group or solvent ions is not included in the fit. Indeed, even when focusing on just the effect of solvent, the IPolQ protein parameterizations suggest that (HF/6-31G*) is a crude representation of this physics, particularly for polarizable groups such as the protein backbone. In contrast, the sSAPT0/jun-cc-pVDZ^57^ used in this work is more physically accurate. In RESP, the ESP is used to fit partial charges; in contrast, we used electrostatic interaction energies as targets, similar to the dihedral-fitting approach^17,29^.

A major drawback of the point-charge representation in this work is that, because neighboring residues to interacting RNA pairs are represented as point charges, dispersion interactions with the QM atoms (i.e., the interacting RNA pair) and with solvent molecules are absent. Stacking interactions between QM atoms and neighboring bases are likely stabilized by favorable dispersion interactions. The interaction of the RNA solute with solvent molecules far from the RNA is likely dominated by electrostatics, captured by the image mesh. In contrast, nearby solvent molecules that are not replaced by image charges are close enough to exert dispersion forces on the RNA atoms. Future work could focus on improving this model by including dispersion interactions from neighboring residues.

In this first iteration of nonbonded parameter fitting, we focus on optimizing parameters for the standard Amber functional form. We use Lorentz-Berthelot^82,83^ mixing rules for LJ parameters, but several alternatives exist, although they are not as commonly used. Subsequent work could modify or expand the Amber functional form. We also use the LJ atom types from Amber ff99, which treat base imine and amino nitrogen as identical^15^. These could be split into distinct atom types. Additionally, other distinct atom types could be introduced for other atoms, with the possibility of tuning individual atoms from nucleobases if a large enough training data set were available.

Another limitation of the current approach is the use of the Amber ff99 parameters as regularization targets. This prevents overfitting in this pilot study, which uses a relatively small set of fitting target conformations. A subsequent expansion of the number of fitting conformations could reduce or eliminate the need for regularization. This would then fit parameters with no dependence on the current ff99^21^ parameters. The dataset we developed here (Table S1) is structurally diverse. It might be useful for other applications requiring an understanding of nonbonded interactions in RNA.

Also, in this study, we used our previously reported method for fitting dihedral potentials via multiple linear regression, using a large set of conformations and QM potentials as targets^29^. We refit the RNA backbone dihedral parameters α, β, γ, ε, ζ, and χ to QM potential energies. We left the other dihedral parameters the same as the Amber ff99 values. In future work, we could expand the Aytenfisu et al.^29^ method to fit additional dihedral parameters, for example to include dihedrals in the sugar ring or to relax planarity to exocyclic amines^84–87^. In the CHARMM force field, a revised dihedral controlling the 2’ OH orientation was found to improve agreement with experiments^88^.

Finally, the RNA.ROC26 force field is appropriate for modeling RNAs composed of the four canonical ribonucleotides. However, covalently-modified RNA residues play key roles in the biological functions of endogenous RNAs and in the pharmacological functions of therapeutic RNAs. While force field parameters are available in the literature for some common RNA residues with covalent modifications^89,90^, it is arduous and human-time-intensive to extend an atom-typed force field to describe new chemistry. An alternative approach for assigning parameters to molecular systems is direct chemical perception based on cheminformatics strings^91^, which has been pioneered by the Open Force Field (OpenFF) Initiative^92^ and used in their general small molecule force field OpenFF Sage^93,94^. The approach described in this work of using SAPT interaction energies with electrostatic embedding to represent local structural and solvent atoms as a training target to intermolecular interactions could also be used to derive nonbonded parameters for an OpenFF force field, permitting coverage of covalently modified RNAs through parameters inherited or refit from a Sage-like general small molecule force field.

## METHODS

### Curating a diverse data set of training structures using clustering

Version 2.217 of the Bowling Green State University non-redundant RNA dataset with a resolution cutoff of 4.0 Å was used for choosing training conformations^47^. Custom scripts were written using the LOOS^95^ package to identify and extract interacting pairs, and Amber LEaP^14^ was used to truncate the 3’ and 5’ ends and add hydrogens.

To cluster potential training conformations, we used a metric that was determined by the Amber force field functional form. Considering eq. 4 as a linear combination of the pairwise parameters (A_αβ_, B_αβ_, C_αβ_),

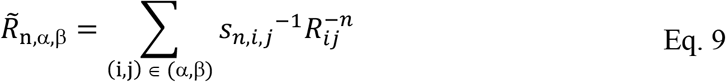

where s is the 1-4 scaling factor for nonbonded interactions. *s*_*12,i,j*_ = *s*_*6,i,j*_ = 2 (LJ) and *s*_*1,i,j*_ = 1.2 (electrostatics) for 1-4 bonded atom pairs.

The nonbonded energy terms in the force field can therefore be written as:

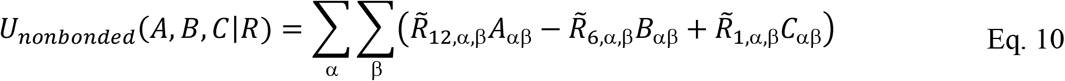

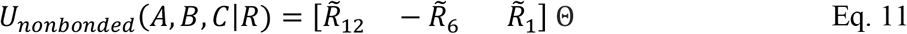

where 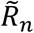 are vectors of the coefficients 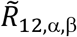 for all pairs of atom types α and β, and Θ is a vector of pairwise parameters, *A*_*α,β*_, *B*_*α,β*_, and *C*_*α,β*_, for all pairs of atom types. For a dataset of M conformations containing *K* LJ atom type pairs and *L* charge type pairs, we create an an *M* × (*2K + L*) matrix:

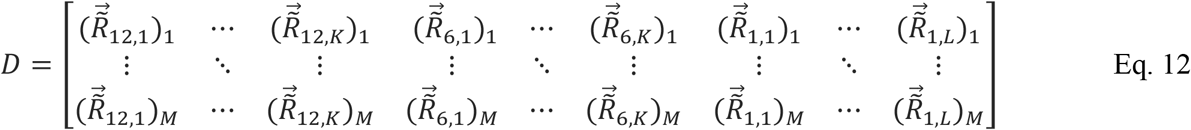

such that:

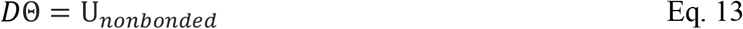

*U*_*nonbonded*_ is then a vector of nonbonded energies for all conformations, *M*. We refer to *D* as the pairwise distance matrix, and each column of this matrix represents the variations in the contribution of one pair of atom types to one of the three terms in the nonbonded term.

The distance metric used for clustering candidate conformations is a standardized Euclidean distance in a space defined by the columns of the pairwise distance matrix, D, which associates pairs of heavy atoms. Hydrogens are added using Amber LEaP^14^ and therefore their positions are not as well determined as heavy atoms. Because of this and to also reduce the dimensionality of the space, hydrogen coordinates were not included in the pairwise distance matrix used for clustering.

In density peaks clustering^50^, cluster centers are areas of high local density surrounded by lower-density regions. Each candidate conformation was assigned a density based on pairwise distances other candidate conformation. A Gaussian kernel was used to estimate the density:

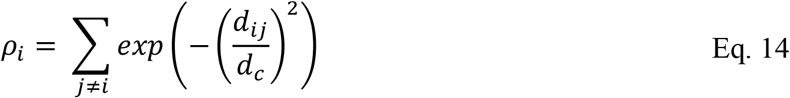

where *d*_*c*_ is the width of the Gaussian kernel equal to the 2^nd^ percentile of the set of all pairwise distances. The minimum distance from any point with higher density was calculated for each point:

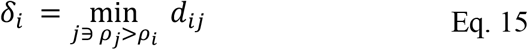

A decision graph was plotted with δ vs ρ (Figure S12). A scoring function was defined to score the likelihood that a point is a cluster center:

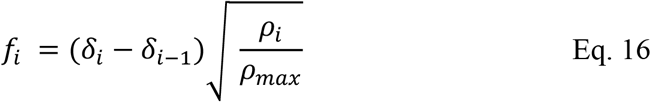

The scoring function assigned higher scores to points with higher values of δ than to points of similar density and was biased towards points with higher density to avoid outliers. The square root function monotonically increases between zero and one, and its first derivative monotonically decreases so that the bias was strongest for points with low density.

### Additional conformations for base-phosphate and base-sugar interactions

Base phosphate and base-ribose sugar interactions were identified from the *Haloarcula marismortui* large ribosomal subunit (PDB ID 1S72)^96^. The list of nucleobase-phosphate and nucleobase-ribose interactions in the 23S rRNA in this structure (chain ID 0) were annotated by FR3D from the BSGU website^4^. Zirbel et al. classified these interactions into 17 types based on nucleobase identity and hydrogen bond donor. For both types of interactions studied here, multiple examples of each of the 17 types of interactions were annotated in chain 0 of 1S72. The distance between the nucleobase heavy atom hydrogen bond donor and either the closest phosphodiester oxygen atom for nucleobase-phosphate interaction or the closest ribose carbon atom for nucleobase-ribose interactions was calculated. For each of the 17 types of interactions, we selected two conformations each: one for which this distance was the smallest, and another for which the distance was closest to the Amber ff99 LJ minimum. The residue donating the hydrogen bond and the two residues flanking the hydrogen bond acceptor (phosphate or sugar) were included in the conformation. To include local RNA structure, adjacent RNA residues that interact with any of these primary residues were included in generating the QM conformation. Residues are considered interacting if either their 3’O3’ - 5’P distance is less than 2.32 Å or if the anisotropic position vector^49^ < √2.5, as mentioned in the curating dataset section in the Results.

### Solvent simulations for training conformations

The Amber ff99 + bsc0 + χ_OL3_ force field^16,17,63^ was used to model the RNA atoms, and monovalent ions were modeled using the Li-Merz 12-6 hydration free energy parameters^14^ for OPC^54^. Following the IPolQ method^46^, we used molecular dynamics simulations to sample configurations of solvent molecules around fixed solute atoms. The solvated conformations were energy-minimized using 4000 steps of steepest descent, followed by 4000 steps of conjugate gradient minimization. The final minimized structures were heated from 5 K to 300 K over 80 ps and then equilibrated at 300 K for 20 ps at constant volume with a Langevin collision frequency of 5 ps^-1^. Followed by equilibration for 200 ps at 300 K with a Langevin collision frequency of 1 ps^-1^ at a constant pressure of 1 atm controlled by a Monte Carlo barostat. The production dynamics were performed at 300K at constant volume for 20 ns. We chose constant volume because a barostat would lead to fluctuations in the electrostatics field. All simulations were performed with periodic boundary conditions using the Particle Mesh Ewald method^97,98^ with a 10 Å direct-space cutoff. A 10 kcal mol^−1^ Å^−2^ Cartesian restraint was applied to the solute in minimization, heating, equilibration and production to preserve the chosen conformation.

### Convergence of solvent MD

The root mean squared error in energy is calculated as the product of the time-averaged solvent ESP and the charge of the RNA atom:

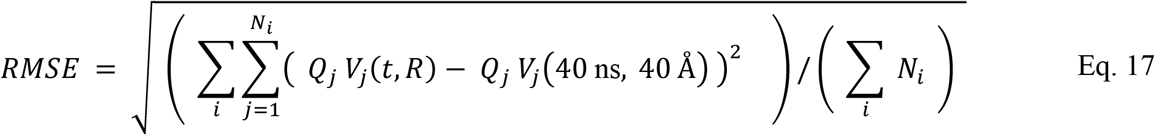

where the sum over *i* runs over conformations in the validation set, the sum *j* over atoms in the conformations, *i*, and *V*_*j*_(*t, R*) is the time-averaged electrostatic potential for the simulation length *t* and solvent cut-off distance *R*. The denominator sums the number of atoms across conformations *i, N*_*i*_.

### Point charges to represent solvent electrostatic field and local environment effects

A mesh was constructed as a union of spherical shells of radius 3 Å centered on the RNA atoms (Figure 4). To develop the mesh points and charges, a set of equidistant points was generated as a union of spherical shells of radius 3 Å around each RNA atom (Figures 3B and S3B) using a Fibonacci spiral^99^, and the points were rotated to a random orientation using quaternions^100,101^ sampled from a random point on a 4D hypersphere^102^. Points within 3 Å of another RNA atom were removed to create the RNA-enclosing mesh. The ESP from solvent molecules and ions at the mesh points, excluding those within the mesh, was calculated. A second mesh with a 4 Å radius was constructed and point charges were placed on it; their magnitudes were fit to reproduce the same ESP at the points on the first mesh. These image charges on the second mesh reproduce approximately the same mean electrostatic field from the far solvent atoms (solvent atoms outside the first mesh). A tool we contributed to the LOOS package^95^, *esp_mesh*, was used to calculate the time-averaged solvent ESP over a trajectory and generate image charges.

### QM interaction energy calculations

The image mesh representing solvent ESP and point charges representing local environment effects were combined along with the interacting residues to build the QM input file (Figure 4). The F/I-SAPT module in Psi4^103^ version 1.5 was used for sSAPT0 and I-SAPT calculations. sSAPT0 exchange scaling was applied to both sSAPT0 and I-SAPT energies.

### Dummy atoms and constraints for parameter fitting

A set of dummy atom types were introduced for atoms in the training conformations that participate in the covalent bond but do not contribute to the interaction energy. The six dummy atoms are marked on Figure S5 (orange). The local electronic structure of these atoms would be perturbed compared to true atoms at these positions (terminal hydroxyl groups and ribose carbons flanking a linked phosphodiester) due to the truncation of the QM system or the localization of electronic orbitals during I-SAPT. The dummy atoms allow these atom type parameters to be fit without influencing the parameters associated with the true atoms. The nonbonded parameters for the nonterminal C5’, C3’ and phosphodiester were informed only from the nucleobase-phosphate interactions in the training conformations. Six dummy charge atom types and three dummy LJ atoms types, one each of carbon, hydrogen and oxygen were used.

Five charge constraints were also applied in the fitting step. The sum of the charges on each of the four nucleosides must be zero. The difference between the total charge of phosphodiester group and the terminal hydroxyl group must be 1 e^-^. The sum of the charges of the dummy atoms must be equal to the sum of the charges for their true terminal hydroxyl atoms. The sum of the dummy charges for C5’ and C3’ must be equal to the sum of the charges for the true C5’ and C3’. During the fit procedure, the hydrogen well depth were found to get stuck at the lower bound, 0 kcal/mol. To rectify this the lower bound of hydrogen well depth was set as Amber ff99^21^ well depth, 0.0157 kcal/mol in the cross validation and basin hopping steps.

### Cross validation and basin hopping to find optimal parameters

Eight subsets of the training dataset were constructed by randomly permuting the indices of the training conformations by type: base-base interactions, base-phosphate interactions, and base-sugar interactions. For each subset, the nonbonded parameters were trained on the other seven subsets using local optimization, initialized from each of ten starting estimates. Initial estimates were generated by sampling a Gaussian random variable centered on the current Amber ff99^21^ nonbonded parameters. The fit parameters from the starting estimate that produced the lowest sum of squared residuals (SSR) were used to calculate the SSR for the remaining subset that were not used to train these parameters. The cross-validation error for a given λ was calculated as the minimum test-set SSR across all eight subsets. For partial charges, a 1D search was performed with *λ*_*q*_ ranging from 10^−2^ to 10^4^, and a grid search over LJ half radius and well depth, with *λ*_*Rmin*_ and *λ*_ϵ_ ranging from 10^0^ to 10^4^ (Figure 5).

Cross-validation was implemented using *scipy*.*optimize*.*least_square*^104^. For the first iteration, the optimal regularization strengths are λ_q_ = 10^2.500^, λ_Rmin_ = 10^1.250^, λ_ϵ_ = 10^0.250^; for the second iteration of the fit, λ_q_ = 10^2.125^, λ_Rmin_ = 10^1.250^, λ_ϵ_ = 10^1.875^. These regularization strengths were used to run the basin-hopping algorithm starting from ten initial estimates. The basin-hopping algorithm was implemented in Python using the *scipy*.*optimize*.*basin_hopping*^104^ function. The temperature parameter for the Metropolis-Hastings criterion in the Monte Carlo optimization was 10 kcal^2^ mol^-2^. The fit partial charges, LJ well depths, and LJ half radii from the two iterations are in Table S4. In the first iteration of charge fit, basin-hopping optimization was run for both minima found in cross-validation (Figure 5), λ_q_=10^1.125^ and λ_q_=10^2.500^, and the parameters from λ=10^1.125^ showed strongly polarized heavy atoms and exhibited greater variation than those from Amber ff99^21^. In the LJ cross-validation for both iterations, the minimum SSR was found in regions where λ < 10^0^. Still, in this region, we hypothesized that the regularization strengths would be too low, leading to overfitting. Hence, the regions with higher regularization strength and minimal SSR were chosen.

Basin-hopping was performed with ten starting estimates generated by sampling a Gaussian distribution, and the parameters were accepted if all ten optimizations converged to the same set of fit parameters and were deemed a global minimum. If not, the next-best regularizations with the lowest SSR were used to run the basin-hopping optimization again. Similar to charge fitting, different regularization strengths with low SSR values were used to run basin hopping.

### Explicit solvent benchmarks

The RNA molecules were solvated in a truncated octahedron of OPC water with 10 Å of padding. K^+^ counterions were added to neutralize the RNA’s charge, and K^+^/Cl^-^ ions were added to achieve a bulk salt concentration of 150 mM^54,55^. The solvated conformations were energy minimized using 4000 steps of steepest descent, followed by 4000 steps of conjugate gradient minimization. The final minimized structures were heated from 5 K to 300 K over 80 ps and then equilibrated at 300 K for 20 ps at constant volume with a Langevin thermostat collision frequency of 5 ps^-1^. Followed by equilibration for 200 ps at 300K with a Langevin thermostat collision frequency of 1 ps^-1^ at a constant pressure of 1 atm controlled by a Monte Carlo barostat. A 10 kcal mol^−1^ Å^−2^ Cartesian restraint to the starting PDB structure was used for minimization, heating and equilibration. The production dynamics were performed at 300 K, at constant pressure of 1 atm, for 2.5 μs for the hairpins and 2 μs for duplexes and tetramers. Simulations were performed with periodic boundary conditions using the Particle Mesh Ewald method with a 10 Å direct-space cutoff^97,98^.

For building tetramer benchmark simulations, the starting structure was generated using NAB^14^ to generate an ideal A-form starting structure. After energy-minimizing each structure, a 200 ps simulated annealing run was performed to generate starting structures for benchmark simulations. The system was heated from 300 K to 500 K in 20 ps, followed by gradual cooling to 300 K over 176 ps and a 4 ps equilibration at 300 K. The other simulation details then were the same as the hairpins and duplexes, except that the direct-space cutoff was 8 Å for tetramers.

The root-mean-square deviation (RMSD) of heavy atoms was calculated for all trajectories relative to the NMR structure deposited in the PDB (Table 1). The tetramer simulations were clustered based on RMSD. Conformations were sampled every 100 ps from the trajectories, and DBSCAN clustering was performed^105^. The minimum distance between points required for forming a cluster was 1.2 Å, and a minimum of 100 points was required to form a cluster. All the analyses were done using the cpptraj module in Amber^14^ and LOOS^95^.

## ACKNOWLEDGEMENTS

This work was supported by a National Institutes of Health Grant, R35GM145283, to DHM.

## Figures and Tables

**Figure 18:**
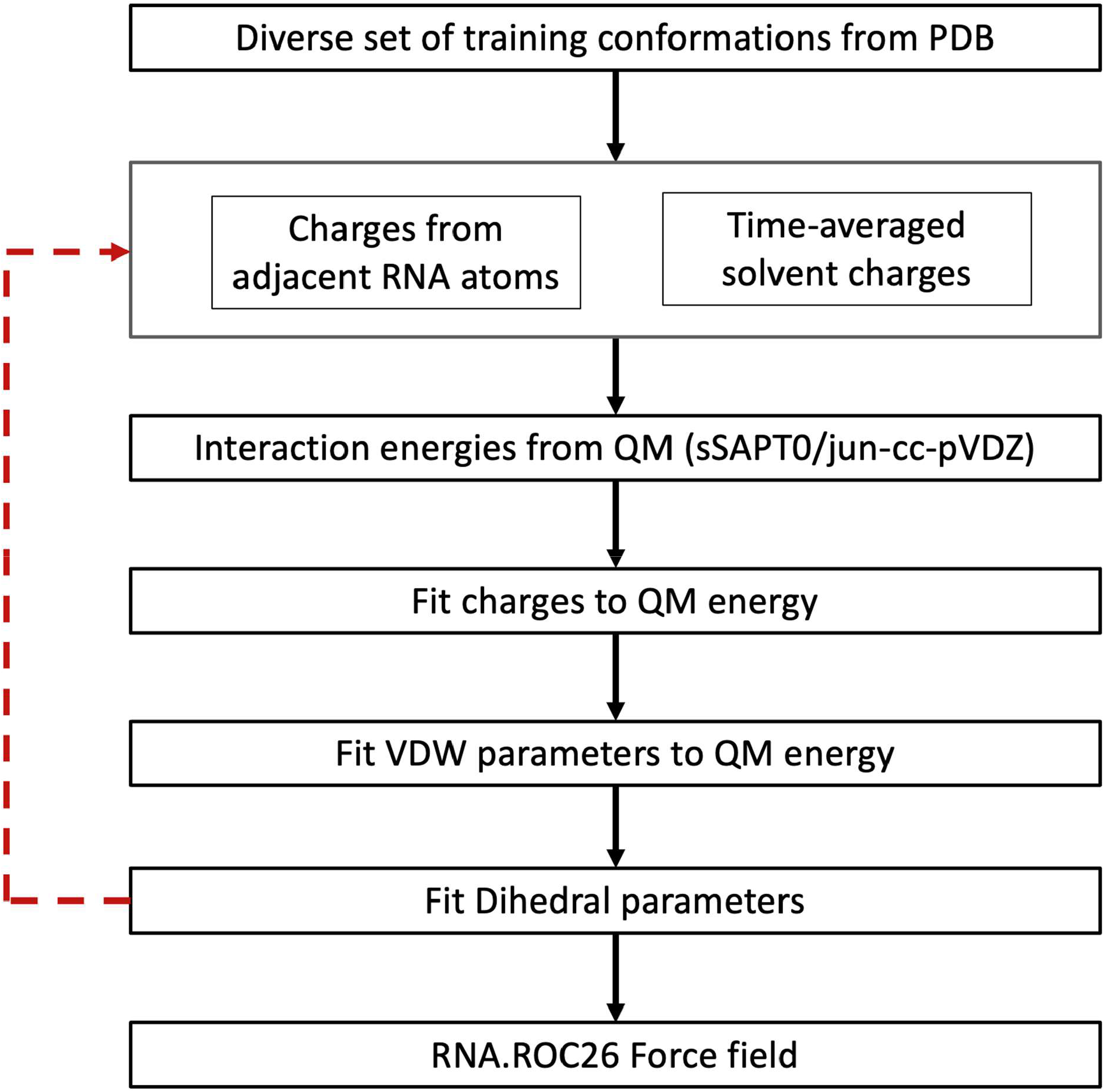
Procedure to fit nonbonded parameters. A diverse set of candidate conformers was assembled. MD simulations were then used to sample the surrounding solvent. Then nonbonded interaction energies were calculated, including solvent charges, using sSAPT0/jun-cc-pVDZ^57^. The interaction energies were used as targets to fit nonbonded parameters, charges first, then VDW parameters, followed by a dihedral parameter fit^29^. This procedure was iterated once back to the MD simulations of the surrounding solvent.

**Figure 19:**
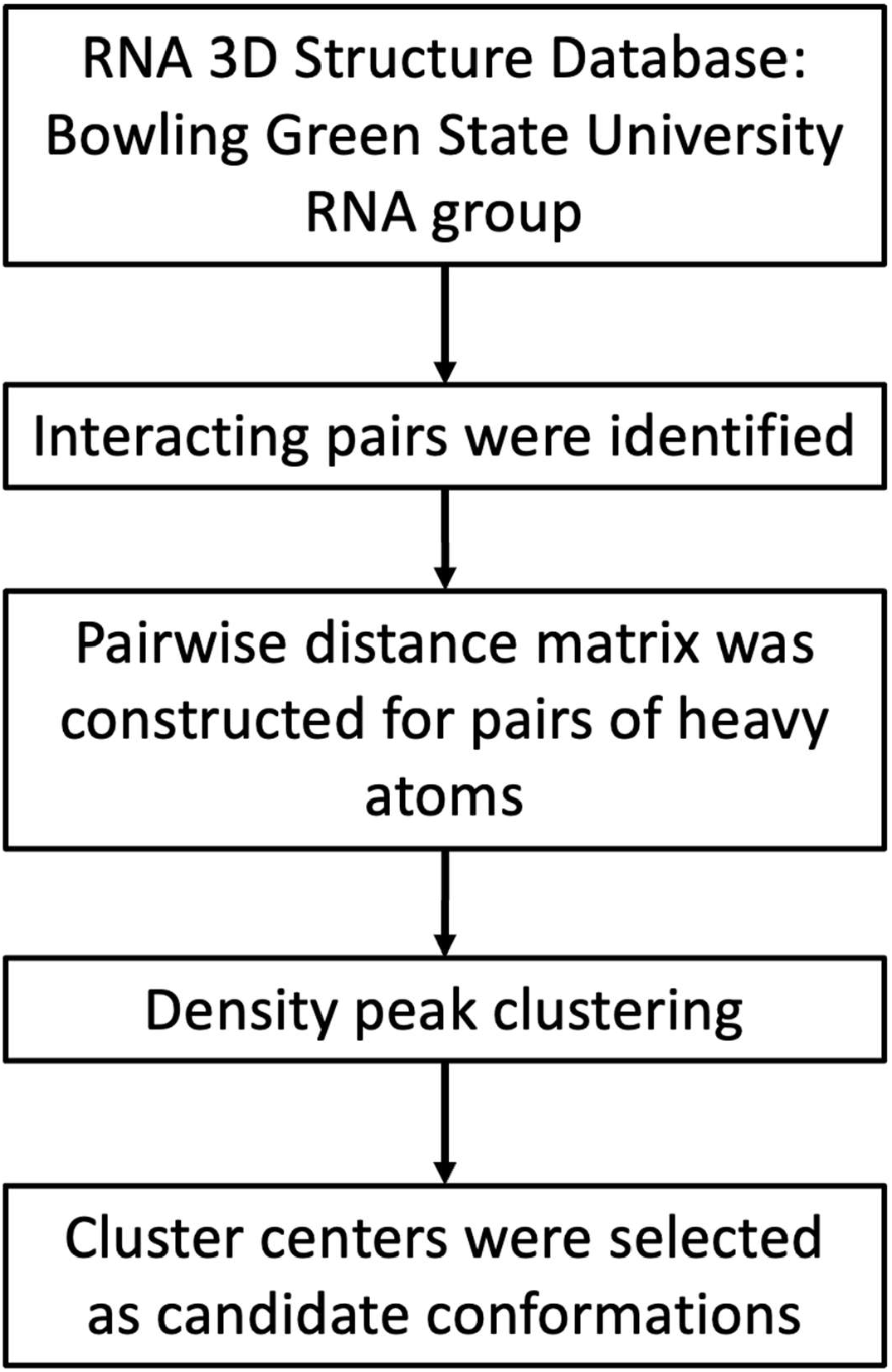
Interacting RNA nucleotide pairs were identified from the RNA 3D structure database^47^, including base stacks and base pairing interactions. A distance metric based on the variations in the nonbonded interaction energy terms due to geometry was used for clustering^106^ these candidate conformations. Structures were clustered by nucleobase identity, and the cluster centers were used as conformations for training nonbonded parameters.

**Figure 20:**
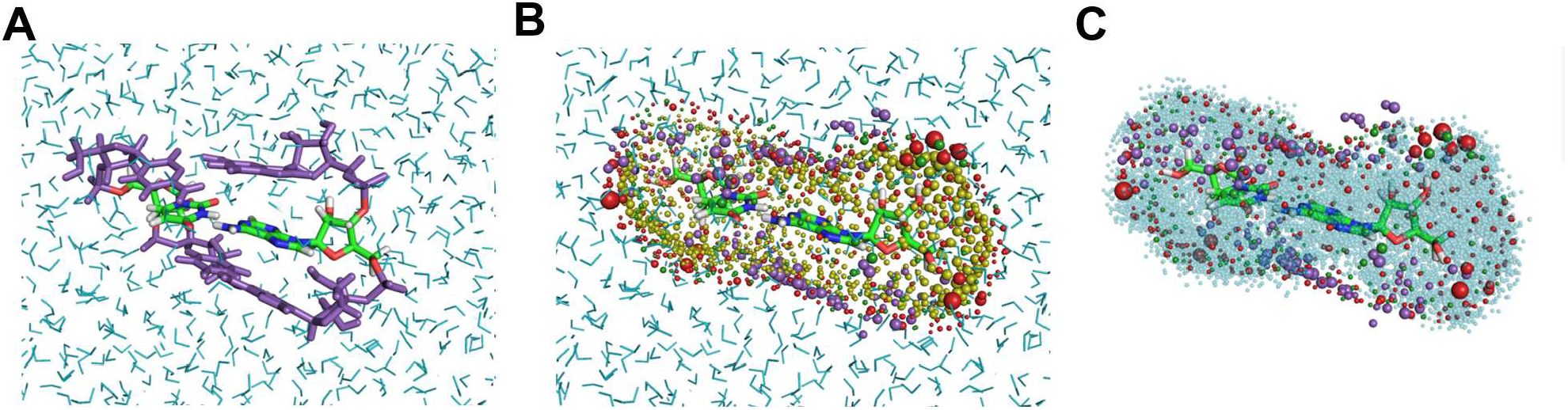
Solvation is included in the training conformations. (A) Equilibrated coordinates of a solvated candidate conformer. The nucleotide pair for QM is shown with atomic coloration, the adjacent neighboring residues are marked in purple, and the solvent molecules are in cyan. (B) Derivation of image charges: yellow spheres represent the mesh, with their sizes indicate the relative magnitude of the solvent ESP. Charges were placed on the image mesh as red spheres for positive and yellow for negative charges. Purple spheres represent neighboring atoms. (C) The final QM input structure after removing the solvent beyond the mesh and using cyan to represent solvent molecules inside the mesh.

**Figure 21:**
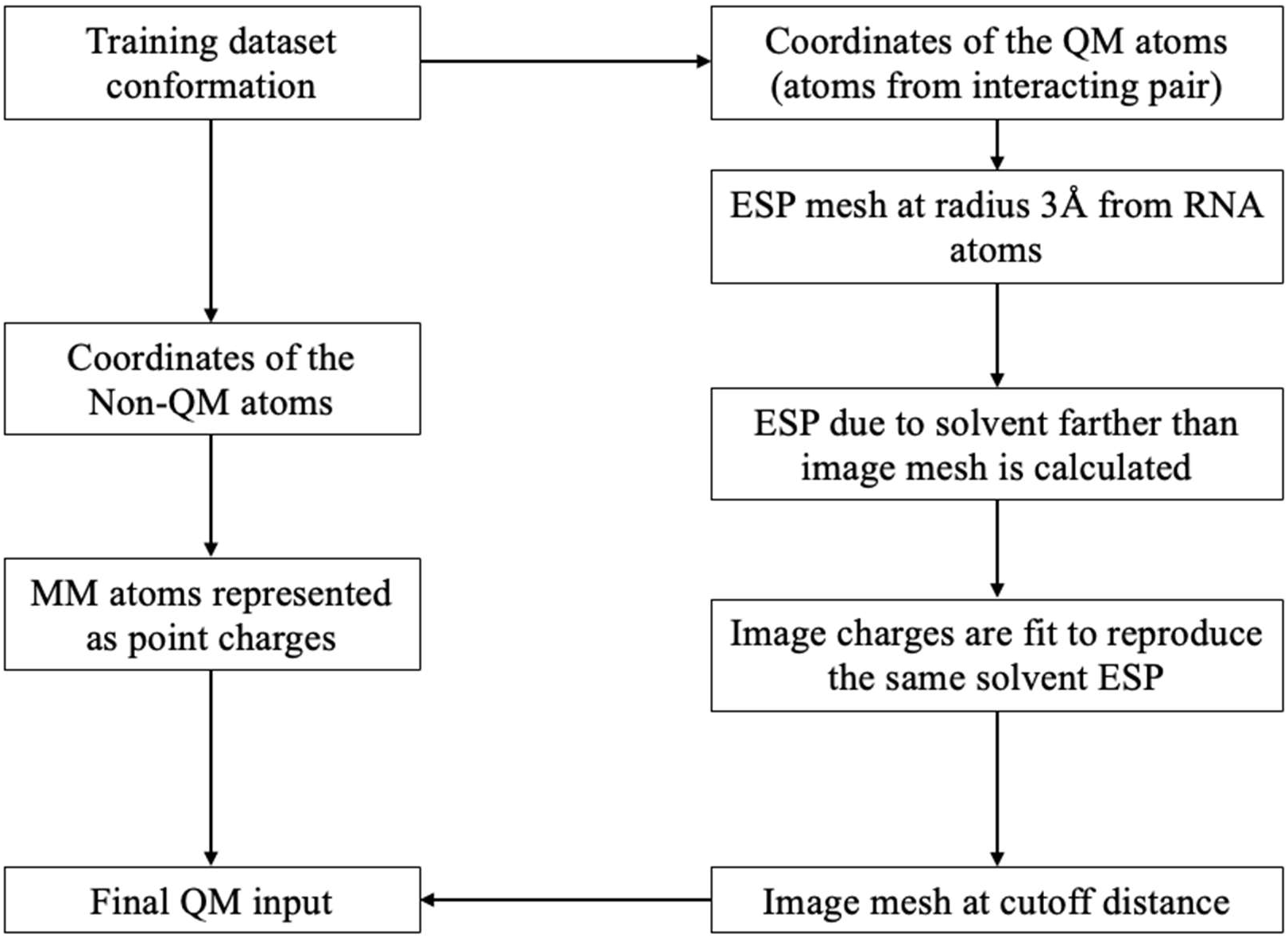
The image charge fitting procedure was used to prepare the input coordinates for the QM calculation. From the input conformation, the downward path indicates that adjacent RNA nucleotides were included as point charges. The path through the right column treated solvent charges to include solvation in the QM. For solvent atoms beyond the mesh, image charges were placed at a cutoff 4 Å distance to reproduce the same ESP as the solvent molecules (illustrated in Figure 3).

**Figure 22:**
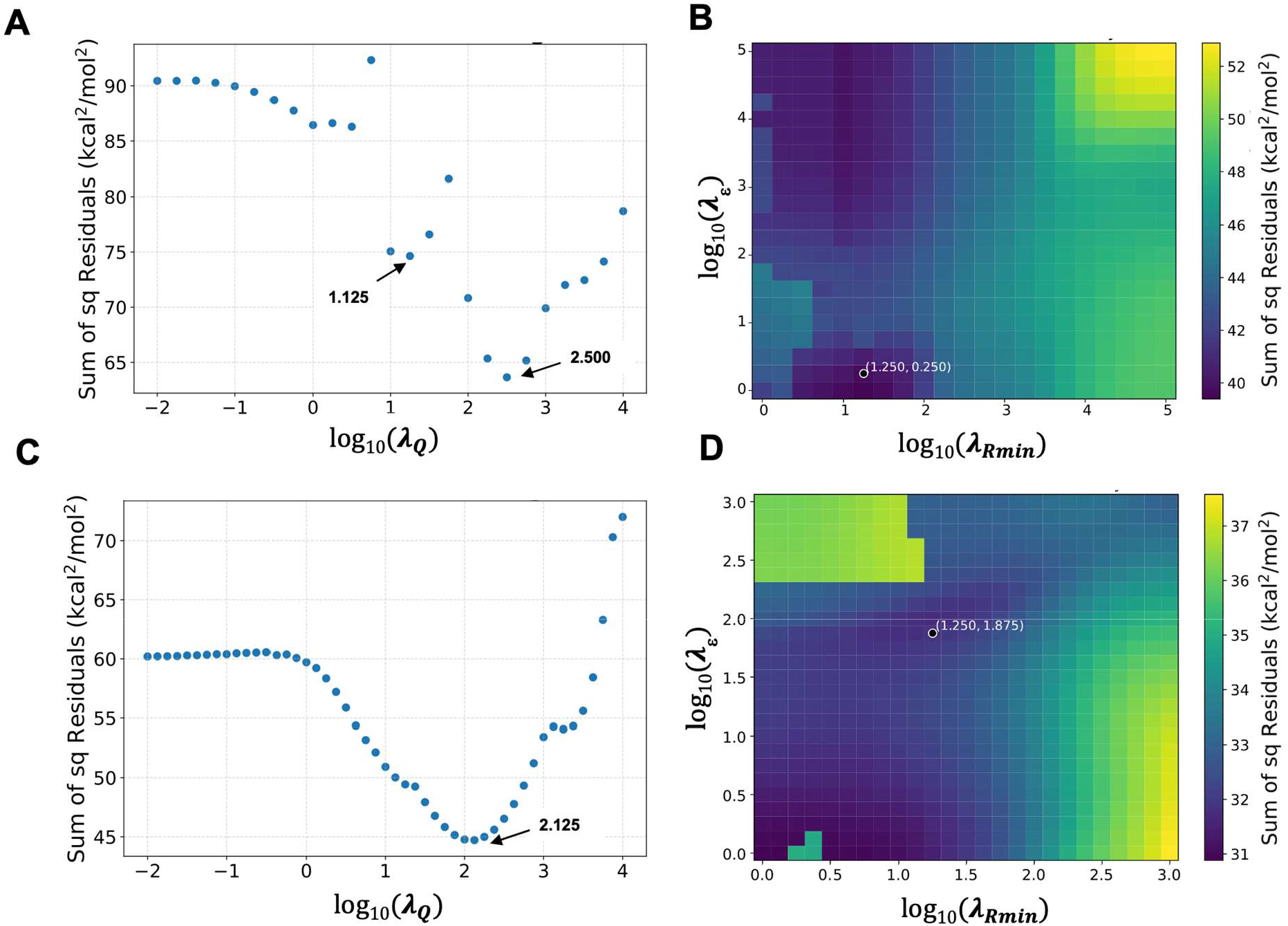
Sum of squared residues (SSR) as a function of regularization strength for charge fits and LJ fits. Charge fit for the first (Panel A) and second (Panel C) iterations. LJ fit for the first (Panel B) and second iteration (Panel D). The chosen regularization constants are illustrated on panels B and D. The training dataset includes 384 conformations and the same training set was used for both iterations of charge and LJ fits.

**Figure 23:**
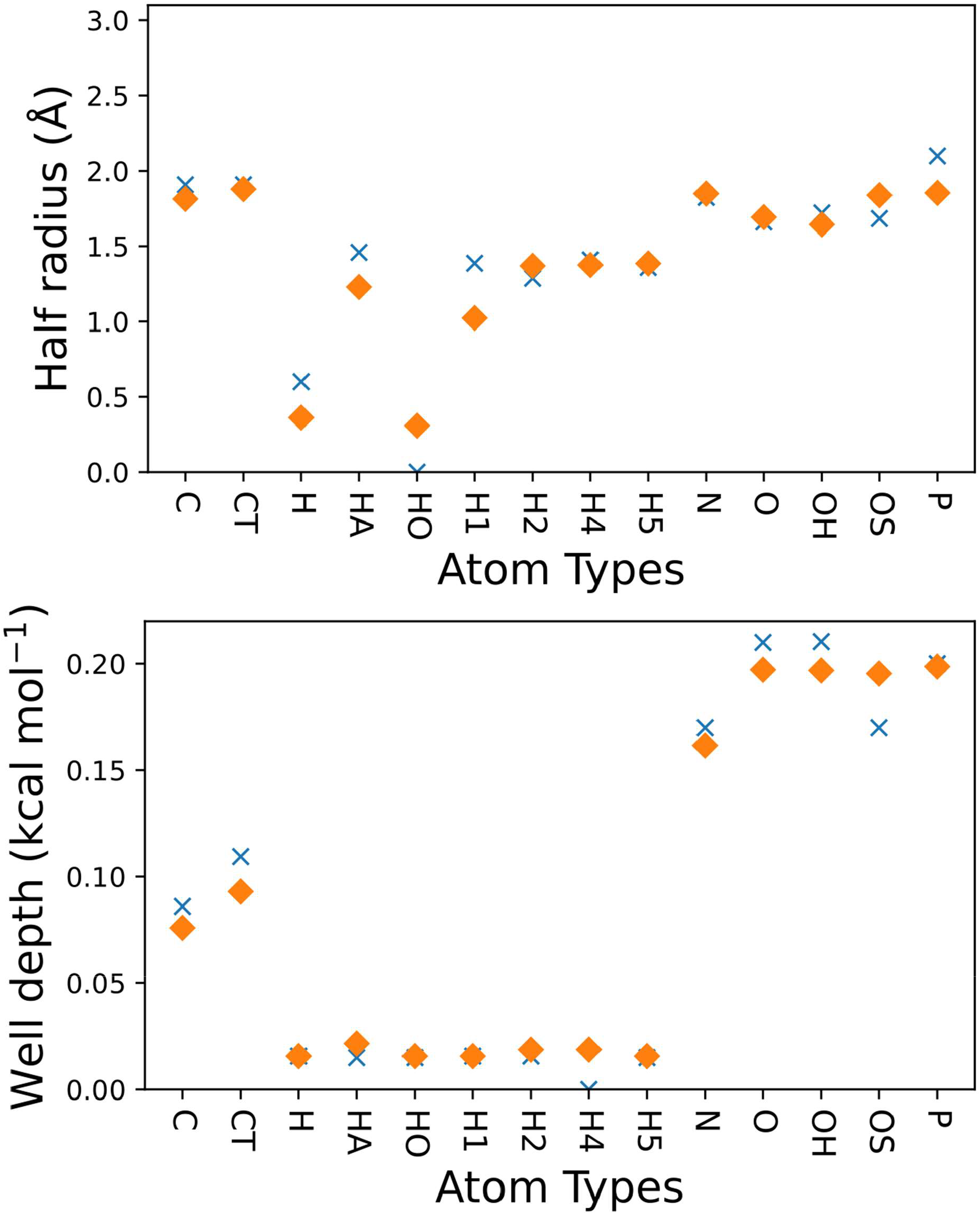
LJ parameters from the fit (orange) and ff99^21^ (blue) compared. Half radius (Panel A) and well depth (Panel B) parameters are plotted.

**Figure 24:**
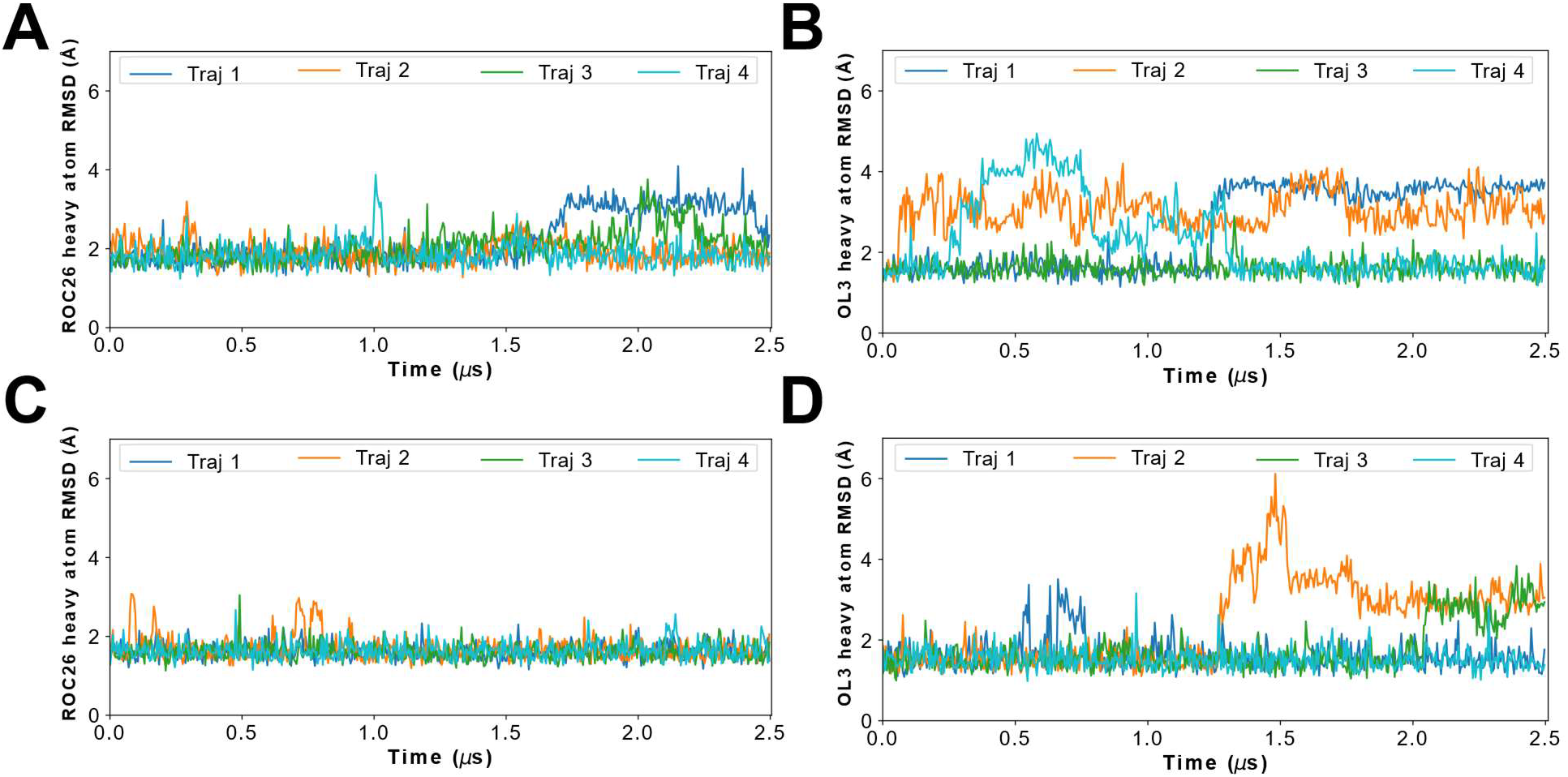
RMSD of heavy atoms as a function of time for 2.5 μs MD simulations of the GCAA tetraloop (PDB ID: 1ZIH; Panel A, B), the UUCG tetraloop (PDB ID: 2KOC; Panel C, D), RNA.ROC26 simulations on the left and RNA.OL3 simulations in the right. The starting NMR structures (Table 1) are used as the reference to calculate RMSD.

**Figure 25:**
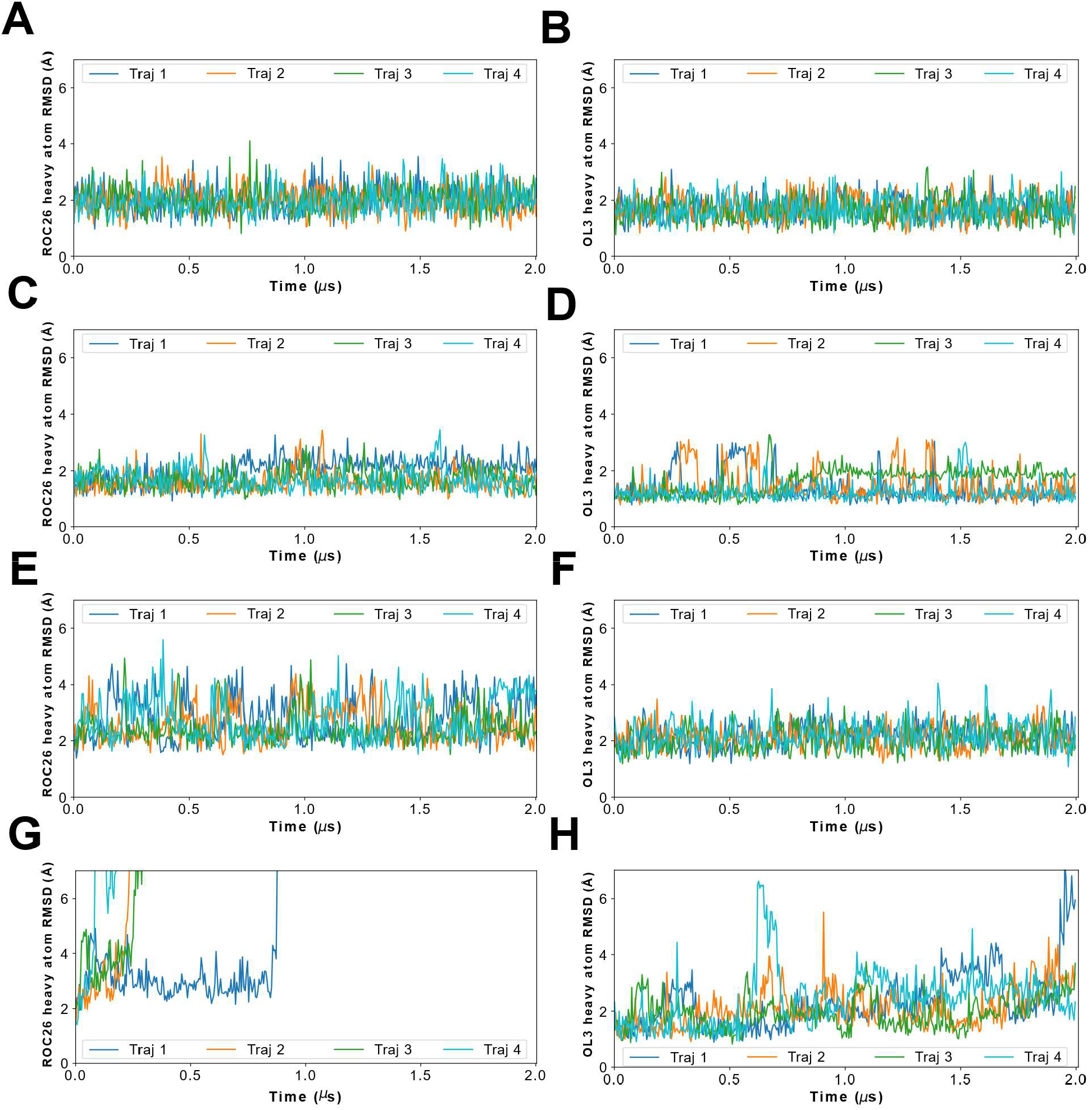
RMSD of heavy atoms as a function of time for 2 μs MD simulations of 2JXQ (A-form duplex; Panel A, B), GC_duplex (Panel C, D), 2DD2 (duplex with an internal loop; Panel E, F) and AU_duplex (Panel G, H). The starting NMR/NAB^14^ structures (Table 1) were used as the reference to calculate heavy atom RMSD.

**Figure 26:**
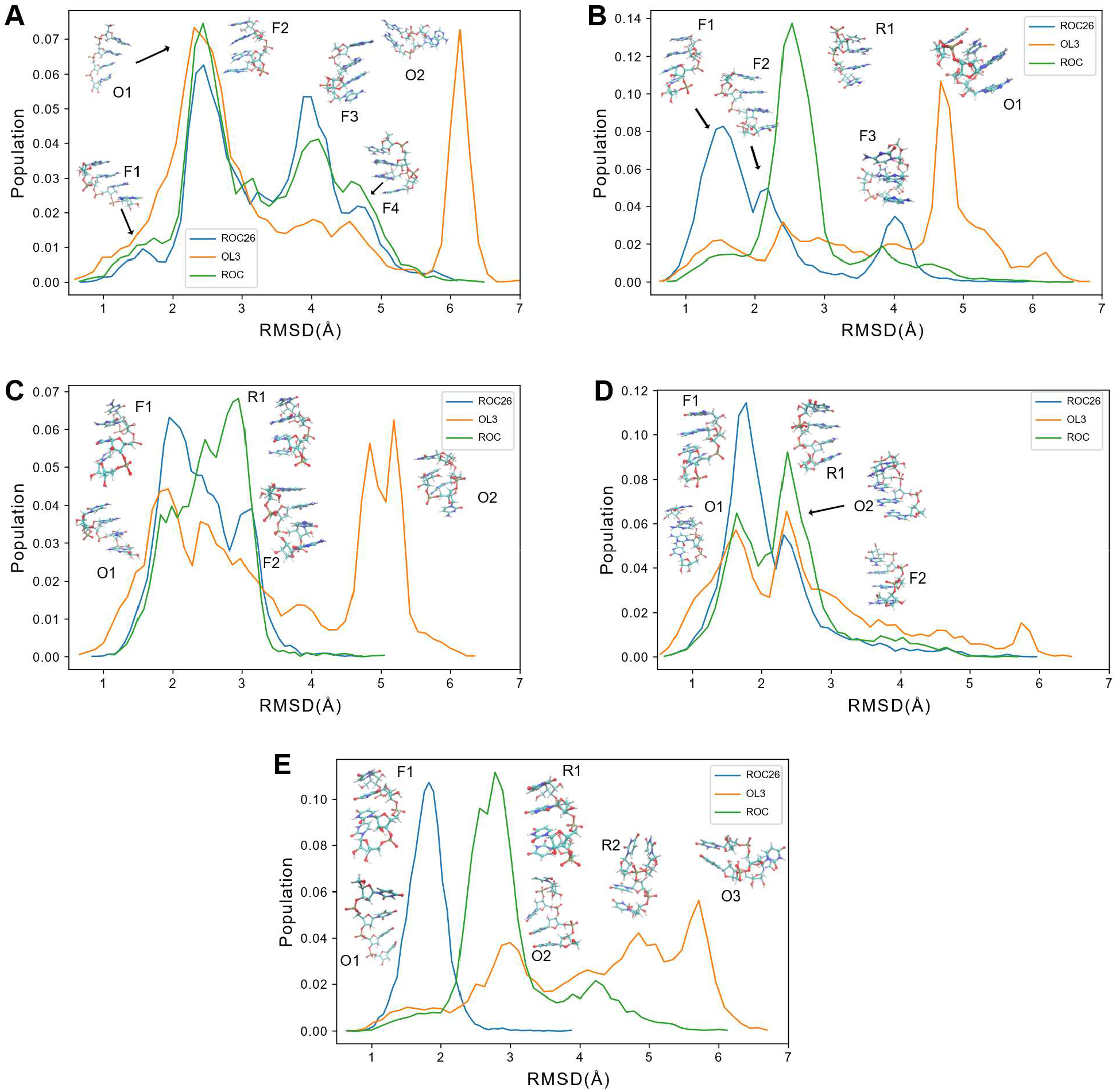
Histogram of root mean squared deviation (RMSD) to A-form starting structure. Panel A shows AAAA, Panel B shows CAAU, Panel C shows CCCC, Panel D shows GACC and Panel E shows UUUU. Five independent simulations were merged for each forcefield: RNA.ROC26 (blue), Amber ff99 + bsc0 + χ_OL3_^17^ (orange), and RNA.ROC^29^ (green). Major cluster centers are marked as F* for RNA.ROC26, O* for Amber ff99 + bsc0 + χ_OL3_^17^ and R* for RNA.ROC^29^.

