## Supplementary Figures and Tables for "Reparameterization of the Amber RNA Force Field Non-Bonded Terms"

Table S1: Details of selected structures (cluster centers)

| PDB ID | CHAIN ID A | RESIDUE ID A | CHAIN ID B | RESIDUE ID B | Number of Nucleotides |
|---|---|---|---|---|---|
| **AA** | | | | | |
| 5MRC | A | 337 | A | 338 | 8 |
| 4V88 | A | 262 | A | 285 | 10 |
| 5FDU | 1 | 344 | 1 | 345 | 11 |
| 3EGZ | B | 33 | B | 49 | 8 |
| 3J7A | A | 1111 | A | 1112 | 7 |
| 4MCF | H | 1 | H | 2 | 5 |
| 3D0U | A | 26 | A | 62 | 6 |
| 5MMM | a | 17 | a | 1029 | 8 |
| 2XGJ | D | 2 | D | 3 | 4 |
| 5T2A | A | 56 | A | 57 | 8 |
| 5MMM | a | 1441 | a | 1442 | 4 |
| 5D0A | F | 5 | F | 9 | 7 |
| 3J9M | A | 730 | A | 731 | 6 |
| 4V8P | F | 603 | F | 604 | 8 |
| 5AJ0 | B | 36 | B | 37 | 4 |
| 2XXA | G | 50 | G | 51 | 5 |
| 3SN2 | B | 26 | B | 27 | 8 |
| 5MRC | a | 823 | a | 842 | 7 |
| 5A2Q | 2 | 726 | 2 | 727 | 7 |
| 1Y26 | X | 52 | X | 53 | 6 |
| 4V8P | F | 306 | F | 307 | 8 |
| 5AJ3 | A | 364 | A | 365 | 8 |
| 5AJ3 | A | 308 | A | 309 | 4 |
| 5MMI | B | 58 | B | 59 | 7 |
| 5T2A | A | 652 | A | 653 | 5 |
| 5MRC | A | 2045 | A | 2046 | 4 |
| 5A2Q | 2 | 638 | 2 | 639 | 6 |
| 4V19 | A | 705 | A | 706 | 8 |
| 5LJ3 | V | 207 | V | 208 | 4 |
| 3J7Y | A | 1159 | A | 1160 | 4 |
| 4V88 | A | 222 | A | 223 | 7 |
| 4V19 | A | 1505 | A | 1506 | 6 |
| **AC** | | | | | |
| 5D0B | F | 5 | F | 13 | 7 |
| 4S3N | C | 22 | C | 23 | 8 |

| | | | | | |
|---|---|---|---|---|---|
| 4Z31 | D | 26 | D | 27 | 7 |
| 5HL7 | X | 238 | X | 645 | 10 |
| 377D | D | 9 | D | 10 | 6 |
| 2ANR | B | 2 | B | 23 | 5 |
| 5FDU | 1 | 1939 | 1 | 2532 | 9 |
| 5V0J | B | 11 | B | 12 | 6 |
| 4GMA | Z | 121 | Z | 122 | 7 |
| 3D2V | A | 4 | A | 77 | 6 |
| 5DHB | C | 3 | C | 4 | 4 |
| 4BTS | D | 1407 | D | 1469 | 10 |
| 1DQH | B | 1 | B | 2 | 5 |
| 4V50 | C | 2 | C | 17 | 5 |
| 4V19 | A | 858 | A | 859 | 9 |
| 5MMI | A | 2115 | A | 2116 | 5 |
| 4YVI | C | 56 | C | 59 | 9 |
| 4V8P | F | 687 | F | 826 | 7 |
| 3-Jan | 4 | 176 | 4 | 177 | 7 |
| 5MMI | A | 2310 | A | 2311 | 4 |
| 2GTT | W | 94 | W | 95 | 4 |
| 4WSM | 3 | 72 | 3 | 73 | 5 |
| 5T2A | A | 195 | A | 196 | 8 |
| 4V88 | A | 512 | A | 540 | 8 |
| 3JAM | 2 | 514 | 2 | 542 | 9 |
| 4V88 | 5 | 163 | 5 | 164 | 6 |
| 3J7P | 5 | 1665 | 5 | 1667 | 6 |
| 5U31 | B | 19 | B | 64 | 6 |
| 3J79 | A | 337 | A | 338 | 10 |
| 3J7Y | A | 4 | A | 133 | 6 |
| 5V0J | A | 9 | B | 11 | 6 |
| 3J79 | A | 3156 | A | 3157 | 7 |
| **AG** | | | | | |
| 4V4Q | C | 767 | C | 803 | 8 |
| 5J7L | D | 2349 | D | 2350 | 10 |
| 3D0U | A | 87 | A | 88 | 7 |
| 5LZS | 5 | 906 | 5 | 1071 | 8 |
| 5FDU | 1 | 2628 | 1 | 2635 | 7 |
| 4V88 | 5 | 2882 | 5 | 2883 | 11 |
| 3ZP8 | A | 1 | A | 2 | 3 |
| 4WRT | V | 13 | V | 14 | 6 |

| | | | | | |
|---|---|---|---|---|---|
| 5UH5 | I | 2 | I | 3 | 3 |
| 2F8K | B | 12 | B | 13 | 8 |
| 5J7L | D | 547 | D | 549 | 5 |
| 5HL7 | X | 912 | X | 921 | 7 |
| 5MMI | A | 773 | A | 774 | 10 |
| 4WZJ | Y | 38 | Y | 67 | 7 |
| 4V8C | C | 18 | C | 69 | 9 |
| 3TD1 | B | 13 | A | 31 | 7 |
| 5AJ0 | 2 | 1911 | 2 | 1917 | 10 |
| 5T5H | A | 981 | A | 983 | 8 |
| 5HL7 | X | 1412 | X | 1414 | 8 |
| 5AJ0 | 2 | 3107 | 2 | 3236 | 7 |
| 3J9M | A | 350 | A | 367 | 10 |
| 5LI0 | a | 55 | a | 379 | 10 |
| 4P3U | A | 35 | A | 37 | 9 |
| 1G1X | D | 7 | D | 18 | 9 |
| 3J9M | A | 767 | A | 773 | 11 |
| 5T2A | 2 | 1519 | 2 | 1570 | 7 |
| 1ZX7 | E | 16 | F | 56 | 8 |
| 1X8W | A | 8 | A | 176 | 10 |
| 5AN9 | N | 278 | N | 289 | 8 |
| 3JCS | 2 | 489 | 2 | 491 | 7 |
| 5MMM | a | 1099 | a | 1100 | 7 |
| 4V9F | 0 | 638 | 0 | 753 | 8 |
| **AU** | | | | | |
| 5LJ3 | Z | 104 | Z | 163 | 6 |
| 5MRC | A | 379 | A | 380 | 8 |
| 4V91 | 1 | 2575 | 1 | 2597 | 6 |
| 5MRC | A | 1959 | A | 1960 | 8 |
| 5LJ3 | Z | 106 | Z | 161 | 5 |
| 5LJ3 | Z | 10 | V | 259 | 5 |
| 1FUF | A | 3 | B | 25 | 6 |
| 3D0U | A | 25 | A | 62 | 7 |
| 4U34 | B | 7 | A | 8 | 4 |
| 4MCF | H | 1 | G | 20 | 4 |
| 4V9F | 0 | 1899 | 0 | 2093 | 10 |
| 4P9R | A | 50 | A | 51 | 7 |
| 4V9F | 0 | 446 | 0 | 447 | 11 |
| 4U34 | B | 6 | B | 7 | 6 |

| | | | | | |
|---|---|---|---|---|---|
| 3J9W | B | 887 | B | 981 | 6 |
| 5LZT | 9 | 592 | 9 | 992 | 6 |
| 4V91 | 1 | 3104 | 1 | 3198 | 6 |
| 5LZT | 9 | 458 | 9 | 476 | 6 |
| 4V9F | 0 | 229 | 0 | 427 | 8 |
| 4BTS | D | 616 | D | 1040 | 8 |
| 3J79 | A | 182 | A | 231 | 6 |
| 4U35 | A | 8 | A | 9 | 5 |
| 5AJ0 | 2 | 3510 | 2 | 3607 | 6 |
| 3JAM | 2 | 695 | 2 | 707 | 6 |
| 5MRC | A | 1064 | A | 1112 | 7 |
| 2A64 | A | 63 | A | 81 | 8 |
| 5MRC | a | 889 | a | 1120 | 7 |
| 3J9W | B | 754 | B | 767 | 6 |
| 3J79 | A | 2516 | A | 2606 | 7 |
| 4V88 | A | 217 | A | 824 | 8 |
| 4YAZ | R | 28 | R | 54 | 9 |
| 3J7P | S | 378 | S | 393 | 7 |
| **CC** | | | | | |
| 4JXZ | B | 64 | B | 65 | 7 |
| 5ED2 | F | 6 | E | 41 | 6 |
| 2R20 | A | 2 | B | 24 | 5 |
| 5FJ0 | C | 18 | C | 19 | 3 |
| 2V7R | B | 8 | B | 9 | 5 |
| 1YZ9 | E | 32 | F | 34 | 5 |
| 5HL7 | X | 2351 | X | 2352 | 7 |
| 5AJ0 | 2 | 3592 | 2 | 3593 | 7 |
| 3AM1 | B | 52 | B | 53 | 7 |
| 5LZS | 5 | 2728 | 5 | 2729 | 7 |
| 4V91 | 4 | 3344 | 4 | 3345 | 6 |
| 5MMM | a | 304 | a | 305 | 7 |
| 5FDU | 1 | 685 | 1 | 686 | 7 |
| 5MMI | A | 2299 | A | 2300 | 7 |
| 4IOA | X | 920 | X | 921 | 7 |
| 4R4V | A | 147 | A | 148 | 7 |
| 5AJ0 | 2 | 2058 | 2 | 2059 | 7 |
| 3SZX | B | 25 | B | 26 | 7 |
| 5U3G | B | 42 | B | 43 | 7 |
| 4WZM | B | 6 | B | 7 | 7 |

| | | | | | |
|---|---|---|---|---|---|
| 4V88 | 5 | 2527 | 5 | 2528 | 8 |
| 4WZQ | C | 13 | C | 14 | 7 |
| 4E7A | P | 4 | T | 8 | 5 |
| 4XWF | A | 44 | A | 45 | 8 |
| 5MRC | A | 129 | A | 130 | 9 |
| 4IOA | X | 2600 | X | 2601 | 8 |
| 4QEI | C | 46 | C | 47 | 8 |
| 4PHY | A | 1 | B | 46 | 4 |
| 1NUJ | G | 10 | G | 11 | 7 |
| 5FDU | 1 | 556 | 1 | 557 | 8 |
| 4M4O | B | 57 | B | 58 | 7 |
| 3AM1 | B | 26 | B | 27 | 7 |
| **CG** | | | | | |
| 5LZS | 5 | 278 | 5 | 297 | 6 |
| 2R20 | A | 1 | B | 24 | 4 |
| 4LX6 | A | 15 | A | 16 | 7 |
| 3BNS | C | 12 | D | 31 | 5 |
| 3C3Z | A | 19 | B | 25 | 5 |
| 2Y8Y | B | 3 | B | 16 | 5 |
| 4JAH | E | 7 | F | 10 | 5 |
| 2Q1O | C | 11 | D | 12 | 4 |
| 2GCS | B | 40 | B | 41 | 4 |
| 5LZS | 5 | 1104 | 5 | 1174 | 7 |
| 4RBY | A | 1 | A | 2 | 3 |
| 3L25 | C | 7 | C | 8 | 6 |
| 1JBR | D | 14 | D | 20 | 8 |
| 2V7R | A | 1 | A | 2 | 5 |
| 1GTN | W | 39 | W | 40 | 3 |
| 5LZS | 5 | 3500 | 5 | 3502 | 5 |
| 1S03 | B | 2 | B | 46 | 6 |
| 4V88 | A | 1161 | A | 1444 | 6 |
| 1Y27 | X | 42 | X | 56 | 6 |
| 4K31 | B | 21 | B | 22 | 6 |
| 4R4V | A | 2 | A | 184 | 6 |
| 2AKE | B | 4 | B | 68 | 6 |
| 1FUF | A | 13 | B | 14 | 3 |
| 5LZS | 5 | 1833 | 5 | 2173 | 8 |
| 3J7P | 5 | 949 | 5 | 978 | 6 |
| 5T5H | A | 26 | A | 52 | 6 |

| | | | | | |
|---|---|---|---|---|---|
| 5FDU | 1 | 2125 | 1 | 2136 | 6 |
| 5LI0 | a | 771 | a | 812 | 8 |
| 3J7P | 5 | 459 | 5 | 575 | 7 |
| 4V4Q | C | 111 | C | 308 | 8 |
| 5GAM | W | 4 | W | 16 | 6 |
| 4YAZ | R | 49 | R | 67 | 7 |
| **CU** | | | | | |
| 5AXM | P | 50 | P | 61 | 6 |
| 5AJ0 | 2 | 1153 | 2 | 1154 | 7 |
| 4WFL | A | 76 | A | 77 | 7 |
| 1XPE | B | 24 | B | 25 | 5 |
| 5D8T | A | 4 | A | 5 | 6 |
| 5J7L | D | 1930 | D | 1945 | 6 |
| 4V91 | 1 | 450 | 1 | 530 | 6 |
| 280D | C | 11 | C | 12 | 6 |
| 5AY3 | B | 15 | B | 16 | 6 |
| 2DLC | Y | 48 | Y | 49 | 5 |
| 4V91 | 1 | 1595 | 1 | 1749 | 6 |
| 5DHB | C | 1 | D | 7 | 5 |
| 205D | A | 4 | A | 5 | 7 |
| 5AN9 | N | 679 | N | 680 | 8 |
| 5T5H | B | 582 | B | 583 | 8 |
| 4V8P | F | 1064 | F | 1065 | 7 |
| 5J7L | D | 2130 | D | 2131 | 7 |
| 4K4V | F | 8 | F | 9 | 4 |
| 4IOA | X | 1315 | X | 1316 | 7 |
| 3J7A | 7 | 48 | 7 | 62 | 6 |
| 1QA6 | D | 54 | D | 55 | 9 |
| 4JGN | E | 10 | E | 11 | 4 |
| 2ZUE | B | 57 | B | 58 | 6 |
| 5LZT | 9 | 1603 | 9 | 1604 | 7 |
| 2OIU | P | 25 | P | 38 | 7 |
| 3JCS | 2 | 229 | 2 | 230 | 9 |
| 5LZS | 5 | 2628 | 5 | 2629 | 8 |
| 5T2A | 2 | 171 | 2 | 172 | 8 |
| 4IOA | X | 212 | X | 213 | 7 |
| 4V91 | 1 | 1629 | 1 | 1630 | 8 |
| 4V9L | A | 55 | A | 62 | 7 |
| 5T2A | 2 | 1504 | 2 | 1505 | 7 |

| | | | | | |
|---|---|---|---|---|---|
| **GG** | | | | | |
| 5LZS | 5 | 473 | 5 | 550 | 8 |
| 4V9B | A | 4 | A | 5 | 7 |
| 5FDU | 1 | 1468 | 1 | 1486 | 9 |
| 4ERD | C | 1 | C | 2 | 6 |
| 3MEI | B | 43 | B | 44 | 6 |
| 3C3Z | A | 18 | B | 25 | 7 |
| 3TD0 | B | 10 | A | 35 | 7 |
| 3J79 | A | 352 | A | 363 | 9 |
| 3J92 | 7 | 60 | 7 | 61 | 8 |
| 1JZV | D | 7 | C | 10 | 7 |
| 5AJ0 | 3 | 3624 | 3 | 3625 | 7 |
| 5LZS | 5 | 610 | 5 | 669 | 11 |
| 5FJ4 | H | 22 | H | 23 | 7 |
| 4IOA | X | 106 | X | 107 | 7 |
| 4R4V | A | 53 | A | 54 | 8 |
| 5MMI | A | 1156 | A | 1157 | 7 |
| 3J92 | 7 | 107 | 7 | 108 | 9 |
| 5J7L | D | 1906 | D | 1907 | 7 |
| 1LNT | A | 1 | B | 21 | 5 |
| 2E9R | B | 11 | B | 12 | 7 |
| 5LZS | 5 | 2055 | 5 | 2056 | 7 |
| 5LZS | 5 | 2794 | 5 | 2795 | 9 |
| 4V4Q | C | 923 | C | 924 | 8 |
| 5HL7 | X | 1560 | X | 1561 | 7 |
| 5J7L | D | 696 | D | 697 | 8 |
| 5E54 | B | 31 | B | 32 | 7 |
| 5T5A | A | 10 | A | 11 | 8 |
| 4BTS | D | 421 | D | 422 | 8 |
| 1YZ9 | C | 11 | F | 44 | 6 |
| 4BTS | D | 555 | D | 556 | 8 |
| 5T5H | B | 251 | B | 252 | 8 |
| 4V4Q | C | 420 | C | 421 | 8 |
| **GU** | | | | | |
| 4V88 | 5 | 2645 | 5 | 2666 | 9 |
| 2ZKO | D | 33 | D | 34 | 8 |
| 3J9W | B | 1949 | B | 1950 | 7 |
| 3K5Q | B | 1 | B | 2 | 3 |
| 2ZI0 | C | 19 | D | 22 | 5 |

| | | | | | |
|---|---|---|---|---|---|
| 1ZX7 | E | 16 | E | 17 | 7 |
| 4NFQ | B | 4 | C | 28 | 5 |
| 5MRC | A | 191 | A | 208 | 8 |
| 1DQH | A | 18 | A | 19 | 6 |
| 5LZT | 9 | 1019 | 9 | 1020 | 7 |
| 5AMR | B | 8 | C | 17 | 5 |
| 5FDU | 1 | 489 | 1 | 490 | 12 |
| 2GDI | Y | 1 | Y | 78 | 4 |
| 5MRC | A | 1310 | A | 1313 | 7 |
| 3J9W | A | 844 | A | 848 | 7 |
| 4V91 | 1 | 1893 | 1 | 1904 | 6 |
| 5A2Q | 2 | 1061 | 2 | 1331 | 8 |
| 5MRC | A | 97 | A | 131 | 8 |
| 3J9W | A | 985 | A | 1224 | 7 |
| 4V88 | 5 | 2228 | 5 | 2720 | 8 |
| 5T5H | B | 743 | B | 832 | 8 |
| 3D0U | A | 37 | A | 51 | 8 |
| 5MRC | b | 18 | b | 55 | 8 |
| 5IMQ | 5 | 18 | 5 | 54 | 10 |
| 4V8P | F | 2428 | F | 2439 | 6 |
| 3J7P | 5 | 2912 | 5 | 2967 | 8 |
| 3J9W | B | 1757 | B | 1773 | 8 |
| 5B2P | B | 81 | B | 84 | 7 |
| 1NTB | A | 17 | B | 32 | 8 |
| 5G2X | A | 472 | A | 515 | 6 |
| 4V8P | F | 1036 | F | 1067 | 6 |
| 5AJ0 | 2 | 3306 | 2 | 3342 | 6 |
| **UU** | | | | | |
| 5MRC | A | 76 | A | 77 | 7 |
| 4QK8 | A | 70 | A | 79 | 7 |
| 4J7L | B | 1 | B | 2 | 3 |
| 2F8K | B | 15 | B | 16 | 6 |
| 4V91 | 1 | 617 | 1 | 734 | 6 |
| 3JCS | 2 | 962 | 2 | 1026 | 6 |
| 5D5L | D | 38 | D | 39 | 7 |
| 4V8P | E | 3258 | E | 3259 | 7 |
| 4IOA | X | 876 | X | 877 | 8 |
| 2OIU | P | 49 | P | 50 | 7 |
| 4V91 | 1 | 3191 | 1 | 3192 | 7 |

| | | | | | |
|---|---|---|---|---|---|
| 5LZT | 9 | 1040 | 9 | 1041 | 7 |
| 5MMI | A | 75 | A | 76 | 7 |
| 5JU8 | A | 60 | A | 61 | 7 |
| 4V91 | 4 | 3324 | 4 | 3325 | 7 |
| 3J79 | A | 583 | A | 584 | 7 |
| 5AJ0 | 2 | 2274 | 2 | 2275 | 8 |
| 5AXM | P | 4 | P | 5 | 8 |
| 3SKI | A | 4 | A | 5 | 7 |
| 4PLX | A | 42 | A | 43 | 7 |
| 4PMW | D | 11 | D | 12 | 4 |
| 5T2A | 2 | 864 | 2 | 865 | 6 |
| 5IT7 | 8 | 66 | 8 | 67 | 6 |
| 5AN9 | N | 900 | N | 901 | 10 |
| 5AN9 | N | 873 | N | 874 | 8 |
| 5MRC | a | 1358 | a | 1359 | 7 |
| 3J79 | A | 1321 | A | 1322 | 7 |
| 5LI0 | a | 1296 | a | 1297 | 7 |
| 4Y1J | A | 97 | A | 98 | 7 |
| 3JAM | 2 | 1376 | 2 | 1377 | 8 |
| 5MRC | A | 1378 | A | 1379 | 10 |
| 4V88 | A | 1284 | A | 1285 | 7 |
| **Base-phosphate interactions** | | | | | |
| 1S72 | 0 | 1648 | 0 | 830 | 11 |
| 1S72 | 0 | 1299 | 0 | 1276 | 9 |
| 1S72 | 0 | 1119 | 0 | 1165 | 12 |
| 1S72 | 0 | 1920 | 0 | 1961 | 14 |
| 1S72 | 0 | 1798 | 0 | 1793 | 8 |
| 1S72 | 0 | 924 | 0 | 991 | 10 |
| 1S72 | 0 | 1968 | 0 | 1799 | 15 |
| 1S72 | 0 | 1876 | 0 | 1879 | 9 |
| 1S72 | 0 | 446 | 0 | 449 | 11 |
| 1S72 | 0 | 1724 | 0 | 1736 | 8 |
| 1S72 | 0 | 1952 | 0 | 1927 | 11 |
| 1S72 | 0 | 2323 | 0 | 2326 | 15 |
| 1S72 | 0 | 1390 | 0 | 1683 | 13 |
| 1S72 | 0 | 1067 | 0 | 1061 | 13 |
| 1S72 | 0 | 1003 | 0 | 614 | 10 |
| 1S72 | 0 | 616 | 0 | 2375 | 12 |
| 1S72 | 0 | 566 | 0 | 569 | 10 |

| | | | | | |
|---|---|---|---|---|---|
| 1S72 | 0 | 1319 | 0 | 1317 | 15 |
| 1S72 | 0 | 1502 | 0 | 1599 | 11 |
| 1S72 | 0 | 873 | 0 | 2060 | 13 |
| 1S72 | 0 | 1142 | 0 | 1151 | 13 |
| 1S72 | 0 | 807 | 0 | 844 | 14 |
| 1S72 | 0 | 828 | 0 | 830 | 9 |
| 1S72 | 0 | 1432 | 0 | 170 | 11 |
| 1S72 | 0 | 910 | 0 | 887 | 13 |
| 1S72 | 0 | 446 | 0 | 448 | 7 |
| 1S72 | 0 | 130 | 0 | 120 | 13 |
| 1S72 | 0 | 1852 | 0 | 1898 | 12 |
| 1S72 | 0 | 424 | 0 | 225 | 9 |
| 1S72 | 0 | 1192 | 0 | 2367 | 13 |
| 1S72 | 0 | 346 | 0 | 283 | 9 |
| 1S72 | 0 | 1286 | 0 | 1289 | 9 |
| **Base-ribose sugar interactions** | | | | | |
| 1S72 | 0 | 1659 | 0 | 1365 | 7 |
| 1S72 | 0 | 2615 | 0 | 2625 | 8 |
| 1S72 | 0 | 2568 | 0 | 2567 | 8 |
| 1S72 | 0 | 2348 | 0 | 2347 | 9 |
| 1S72 | 0 | 902 | 0 | 1002 | 11 |
| 1S72 | 0 | 1584 | 0 | 1529 | 7 |
| 1S72 | 0 | 184 | 0 | 415 | 11 |
| 1S72 | 0 | 2103 | 0 | 2088 | 9 |
| 1S72 | 0 | 472 | 0 | 495 | 12 |
| 1S72 | 0 | 1801 | 0 | 1800 | 8 |
| 1S72 | 0 | 2645 | 0 | 2509 | 8 |
| 1S72 | 0 | 2597 | 0 | 2558 | 11 |
| 1S72 | 0 | 1916 | 0 | 1904 | 11 |
| 1S72 | 0 | 2698 | 0 | 2564 | 11 |
| 1S72 | 0 | 1299 | 0 | 1275 | 7 |
| 1S72 | 0 | 1247 | 0 | 1245 | 8 |
| 1S72 | 0 | 2738 | 0 | 2737 | 9 |
| 1S72 | 0 | 1293 | 0 | 1292 | 7 |
| 1S72 | 0 | 2688 | 0 | 2687 | 10 |
| 1S72 | 0 | 237 | 0 | 235 | 8 |
| 1S72 | 0 | 2634 | 0 | 2417 | 12 |
| 1S72 | 0 | 1813 | 0 | 1832 | 6 |
| 1S72 | 0 | 1444 | 0 | 772 | 12 |

| | | | | | |
|---|---|---|---|---|---|
| 1S72 | 0 | 1336 | 0 | 1335 | 7 |
| 1S72 | 0 | 1173 | 0 | 1110 | 7 |
| 1S72 | 0 | 2298 | 0 | 2297 | 7 |
| 1S72 | 0 | 1832 | 0 | 1814 | 9 |
| 1S72 | 0 | 430 | 0 | 429 | 10 |
| 1S72 | 0 | 1646 | 0 | 1650 | 8 |
| 1S72 | 0 | 2669 | 0 | 2688 | 12 |
| 1S72 | 0 | 2582 | 0 | 1337 | 6 |
| 1S72 | 0 | 948 | 0 | 945 | 9 |

Table 2: Details of selected structures in validation set conformations

| PDB ID | CHAIN ID A | RESIDUE ID A | CHAIN ID B | RESIDUE ID B | Number of Residues |
|---|---|---|---|---|---|
| 5MRC | A | 337 | A | 338 | 8 |
| 3EGZ | B | 33 | B | 49 | 8 |
| 5D0B | F | 5 | F | 13 | 7 |
| 4S3N | C | 22 | C | 23 | 8 |
| 4Z31 | D | 26 | D | 27 | 7 |
| 3D0U | A | 87 | A | 88 | 7 |
| 5LZS | 5 | 906 | 5 | 1071 | 8 |
| 5LJ3 | Z | 104 | Z | 163 | 6 |
| 5MRC | A | 379 | A | 380 | 8 |
| 4V91 | 1 | 2575 | 1 | 2597 | 6 |
| 5MRC | A | 1959 | A | 1960 | 8 |
| 4JXZ | B | 64 | B | 65 | 7 |
| 5ED2 | F | 6 | E | 41 | 6 |
| 2R20 | A | 2 | B | 24 | 5 |
| 5FJ0 | C | 18 | C | 19 | 3 |
| 5LZS | 5 | 278 | 5 | 297 | 6 |
| 2R20 | A | 1 | B | 24 | 4 |
| 4LX6 | A | 15 | A | 16 | 7 |
| 3BNS | C | 12 | D | 31 | 5 |
| 5AXM | P | 50 | P | 61 | 6 |
| 5AJ0 | 2 | 1153 | 2 | 1154 | 7 |
| 4WFL | A | 76 | A | 77 | 7 |
| 1XPE | B | 24 | B | 25 | 5 |
| 4V9B | A | 4 | A | 5 | 7 |
| 4ERD | C | 1 | C | 2 | 6 |
| 2ZKO | D | 33 | D | 34 | 8 |
| 3J9W | B | 1949 | B | 1950 | 7 |
| 3K5Q | B | 1 | B | 2 | 3 |
| 5MRC | A | 76 | A | 77 | 7 |
| 4QK8 | A | 70 | A | 79 | 7 |
| 4J7L | B | 1 | B | 2 | 3 |
| 2F8K | B | 15 | B | 16 | 6 |

Table S3: Regularization targets for LJ parameter fit.

| Element | $R_{min}$ (Å) | $\epsilon$ (kcal mol$^{-1}$) |
|---|---|---|
| C | 1.9080 | 0.0977 |
| O | 1.6886 | 0.1968 |
| N | 1.8240 | 0.1700 |
| P | 2.1000 | 0.2000 |
| Polar H | 0.6000 | 0.0157 |
| Nonpolar H | 1.3802 | 0.0153 |

Table S4: Fit parameters from first and second iterations of the fit. The second iteration was used for the benchmark simulations. Atom names that start with D represent the dummy atoms.

| Type | Atom | FF99 | Fit iteration 1 | Fit iteration 2 |
|---|---|---|---|---|
| **R*** | C | 1.90800 | 1.812707 | 1.81385 |
| **R*** | CT | 1.90800 | 2.115748 | 1.87896 |
| **R*** | H | 0.60000 | 0.342667 | 0.36205 |
| **R*** | HA | 1.45900 | 1.144264 | 1.22913 |
| **R*** | HO | 0.60000 | 0.245021 | 0.30827 |
| **R*** | H1 | 1.38700 | 1.033491 | 1.02317 |
| **R*** | H2 | 1.28700 | 1.246148 | 1.36989 |
| **R*** | H4 | 1.40900 | 1.302821 | 1.37517 |
| **R*** | H5 | 1.35900 | 1.338349 | 1.38547 |
| **R*** | N | 1.82400 | 1.817833 | 1.84922 |
| **R*** | O | 1.66120 | 1.660980 | 1.69296 |
| **R*** | OH | 1.72100 | 1.648867 | 1.64433 |
| **R*** | OS | 1.68370 | 1.713550 | 1.83851 |
| **R*** | P | 2.10000 | 2.444805 | 1.85378 |
| **R*** | DC | 1.90800 | 2.018476 | 2.13537 |
| **R*** | DH | 0.60000 | 0.615940 | 0.61242 |
| **R*** | DO | 1.72100 | 2.018913 | 2.08263 |
| **Eps** | C | 0.08600 | 0.041268 | 0.07595 |
| **Eps** | CT | 0.10940 | 0.049631 | 0.09296 |
| **Eps** | H | 0.01570 | 0.015700 | 0.01570 |
| **Eps** | HA | 0.01500 | 0.121507 | 0.02162 |
| **Eps** | HO | 0.01500 | 0.057460 | 0.01570 |
| **Eps** | H1 | 0.01570 | 0.015700 | 0.01570 |
| **Eps** | H2 | 0.01570 | 0.021544 | 0.01874 |
| **Eps** | H4 | 0.01500 | 0.023450 | 0.01878 |
| **Eps** | H5 | 0.01500 | 0.024384 | 0.01570 |
| **Eps** | N | 0.17000 | 0.236271 | 0.16158 |
| **Eps** | O | 0.21000 | 0.246544 | 0.19713 |
| **Eps** | OH | 0.21040 | 0.186054 | 0.19681 |
| **Eps** | OS | 0.17000 | 0.173455 | 0.19535 |
| **Eps** | P | 0.20000 | 0.171256 | 0.19863 |
| **Eps** | DC | 0.10940 | 0.226765 | 0.10554 |
| **Eps** | DH | 0.01500 | 0.027179 | 0.01605 |
| **Eps** | DO | 0.21040 | 0.248440 | 0.19829 |

| | | | | |
|---|---|---|---|---|
| Q | C2' | 0.06700 | 0.138709 | 0.13195 |
| Q | C3' | 0.20220 | 0.346274 | 0.35300 |
| Q | C4' | 0.10650 | 0.165883 | 0.15387 |
| Q | C5' | 0.05580 | 0.080048 | 0.05754 |
| Q | H2' | 0.09720 | 0.009898 | 0.03291 |
| Q | H3' | 0.06150 | -0.073490 | -0.06679 |
| Q | H4' | 0.11740 | 0.108267 | 0.11765 |
| Q | H5' | 0.06790 | 0.069172 | 0.05205 |
| Q | HO2' | 0.41860 | 0.397315 | 0.38627 |
| Q | O2' | -0.61390 | -0.479982 | -0.46083 |
| Q | O3' | -0.52460 | -0.490191 | -0.48013 |
| Q | O4' | -0.35480 | -0.297841 | -0.26707 |
| Q | O5' | -0.49890 | -0.381001 | -0.42253 |
| Q | OP | -0.77600 | -0.924725 | -0.91467 |
| Q | P | 1.16620 | 1.180841 | 1.17815 |
| Q | HO3' | 0.43760 | 0.371557 | 0.38286 |
| Q | HO5' | 0.42950 | 0.417757 | 0.42105 |
| Q | TO3' | -0.65410 | -0.692076 | -0.70044 |
| Q | TO5' | -0.62230 | -0.637041 | -0.65731 |
| Q | C1'_A | 0.03940 | -0.003772 | -0.02321 |
| Q | C2_A | 0.58750 | 0.595775 | 0.58361 |
| Q | C4_A | 0.30530 | 0.228873 | 0.22364 |
| Q | C5_A | 0.05150 | 0.037472 | 0.02916 |
| Q | C6_A | 0.70090 | 0.702077 | 0.71191 |
| Q | C8_A | 0.20060 | 0.174915 | 0.17464 |
| Q | H1'_A | 0.20070 | 0.276321 | 0.29900 |
| Q | H2_A | 0.04730 | 0.117318 | 0.12163 |
| Q | H6_A | 0.41150 | 0.437459 | 0.42319 |
| Q | H8_A | 0.15530 | 0.136152 | 0.12305 |
| Q | N1_A | -0.76150 | -0.654980 | -0.67843 |
| Q | N3_A | -0.69970 | -0.846854 | -0.84003 |
| Q | N6_A | -0.90190 | -0.960314 | -0.90887 |
| Q | N7_A | -0.60730 | -0.501367 | -0.49471 |
| Q | N9_A | -0.02510 | -0.170159 | -0.15650 |
| Q | C1'_C | 0.00660 | 0.017190 | 0.05320 |
| Q | C2_C | 0.75380 | 0.803370 | 0.77312 |
| Q | C4_C | 0.81850 | 0.810870 | 0.82070 |
| Q | C5_C | -0.52150 | -0.565075 | -0.57188 |
| Q | C6_C | 0.00530 | 0.033498 | 0.02396 |

| | | | | |
|---|---|---|---|---|
| Q | H1'_C | 0.20290 | 0.096253 | 0.09581 |
| Q | H4_C | 0.42340 | 0.464141 | 0.45131 |
| Q | H5_C | 0.19280 | 0.162337 | 0.15517 |
| Q | H6_C | 0.19580 | 0.130018 | 0.14409 |
| Q | N1_C | -0.04840 | 0.021076 | 0.00969 |
| Q | N3_C | -0.75840 | -0.693293 | -0.69524 |
| Q | N4_C | -0.95300 | -1.000000 | -0.96688 |
| Q | O2_C | -0.62520 | -0.738149 | -0.73313 |
| Q | C1'_G | 0.01910 | -0.015805 | -0.01247 |
| Q | C2_G | 0.76570 | 0.700045 | 0.72966 |
| Q | C4_G | 0.12220 | 0.117443 | 0.11461 |
| Q | C5_G | 0.17440 | 0.198104 | 0.20212 |
| Q | C6_G | 0.47700 | 0.542959 | 0.52941 |
| Q | C8_G | 0.13740 | 0.145920 | 0.16929 |
| Q | H1_G | 0.34240 | 0.397506 | 0.39059 |
| Q | H1'_G | 0.20060 | 0.147632 | 0.11268 |
| Q | H2_G | 0.43640 | 0.454456 | 0.43983 |
| Q | H8_G | 0.16400 | 0.130226 | 0.12951 |
| Q | N1_G | -0.47870 | -0.461908 | -0.45799 |
| Q | N2_G | -0.96720 | -0.984015 | -0.96112 |
| Q | N3_G | -0.63230 | -0.683948 | -0.69277 |
| Q | N7_G | -0.57090 | -0.541533 | -0.53946 |
| Q | N9_G | 0.04920 | 0.021292 | 0.03220 |
| Q | O6_G | -0.55970 | -0.616454 | -0.61468 |
| Q | C1'_U | 0.06740 | 0.021969 | 0.02885 |
| Q | C2_U | 0.46870 | 0.481456 | 0.45093 |
| Q | C4_U | 0.59520 | 0.558687 | 0.55875 |
| Q | C5_U | -0.36350 | -0.344178 | -0.36265 |
| Q | C6_U | -0.11260 | -0.132280 | -0.13909 |
| Q | H1'_U | 0.18240 | 0.137991 | 0.15354 |
| Q | H3_U | 0.31540 | 0.415232 | 0.42223 |
| Q | H5_U | 0.18110 | 0.310286 | 0.30412 |
| Q | H6_U | 0.21880 | 0.072929 | 0.11616 |
| Q | N1_U | 0.04180 | -0.012565 | -0.01759 |
| Q | N3_U | -0.35490 | -0.329322 | -0.35546 |
| Q | O2_U | -0.54770 | -0.541719 | -0.53974 |
| Q | O4_U | -0.57610 | -0.632110 | -0.60879 |
| Q | DC3' | 0.20220 | 0.207272 | 0.20370 |
| Q | DC5' | 0.05580 | 0.219050 | 0.20684 |

| | | | | |
|---|---|---|---|---|
| Q | DH3' | 0.43760 | 0.372561 | 0.39335 |
| Q | DH5' | 0.42950 | 0.226090 | 0.27422 |
| Q | DO3' | -0.65410 | -0.675657 | -0.69047 |
| Q | DO5' | -0.62230 | -0.462796 | -0.53094 |

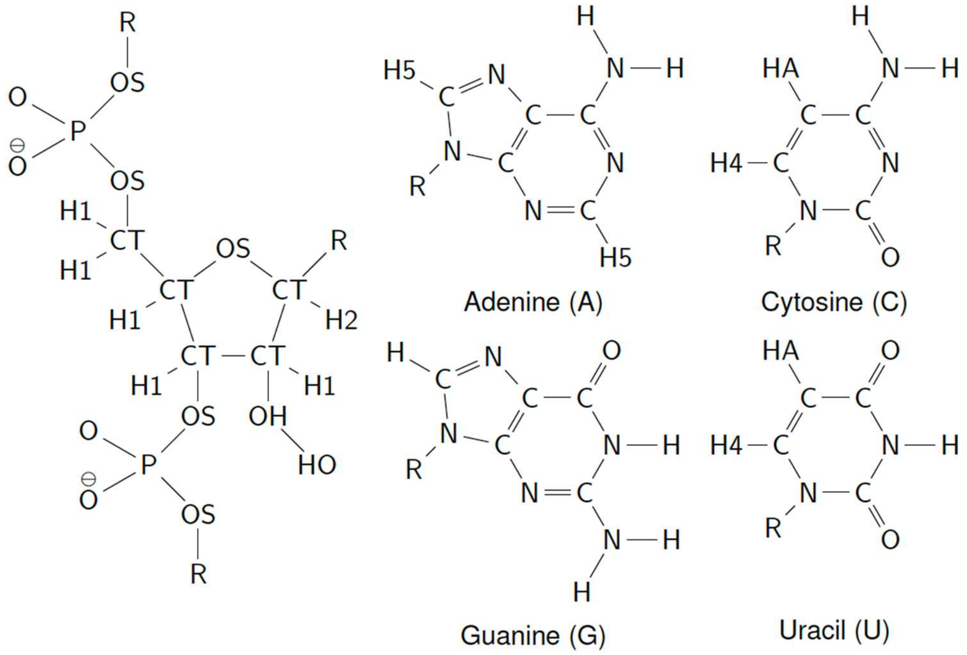


Figure S1: Amber fixed-charge force field LJ atom types for RNA. The sugar phosphate backbone is represented on the left side and four nucleobases: Adenine, Cytosine, Guanine, and Uracil.

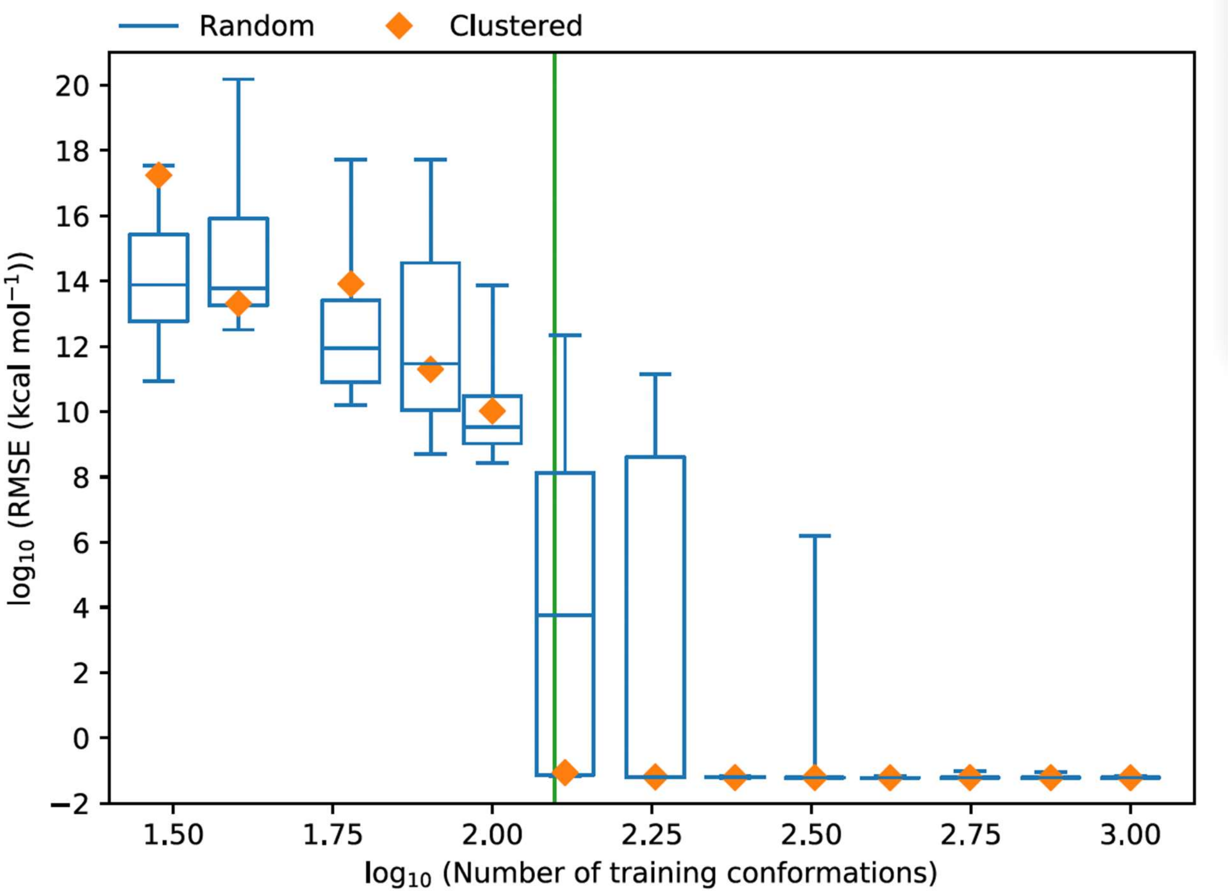


Figure S2: Fits of nonbonded parameters targeting Amber ff99 interaction energies. The root mean square error (RMSE) of the target energies for a test set is plotted against the size of the training dataset. Training datasets constructed by random sampling without replacement are shown as blue box-and-whisker plots; the central line represents the median, the boxes represent the interquartile range, and the whiskers represent the full range across fits to ten random training datasets. These random datasets are a negative control. Clustered datasets constructed from the density peaks clustering method are shown as orange diamonds. The vertical green line represents the number of free parameters in the fit.

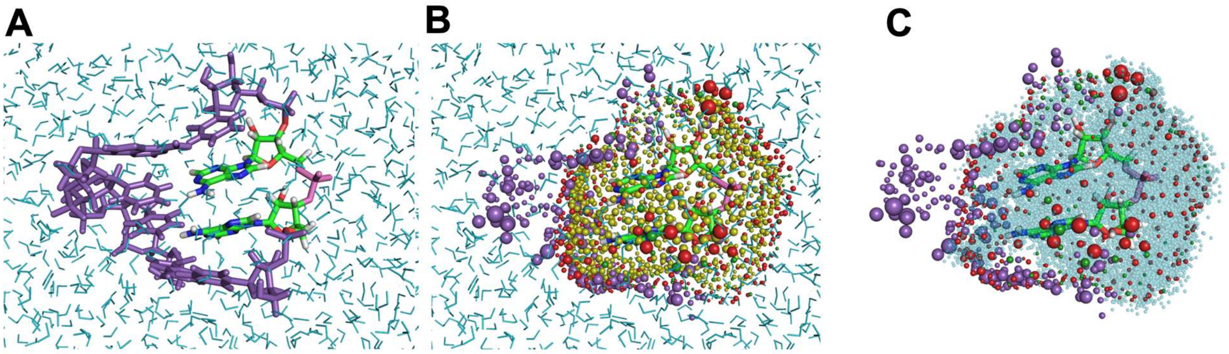


Figure S3: The solvated conformations used for fitting. (A) Minimized coordinates of a solvated candidate conformer with the adjacent interacting nucleotide pair marked in atom coloring, neighboring residues marked in purple, and solvent molecules marked in cyan. (B) Derivation of image charges: yellow spheres represent the mesh, with their sizes indicating the relative magnitude of the solvent ESP. Charges are placed on the image mesh as red spheres for positive and yellow for negative charges. Purple spheres represent neighboring atoms. (C) The final QM input structure after removing the solvent beyond the mesh and using cyan to represent solvent molecules inside the mesh.

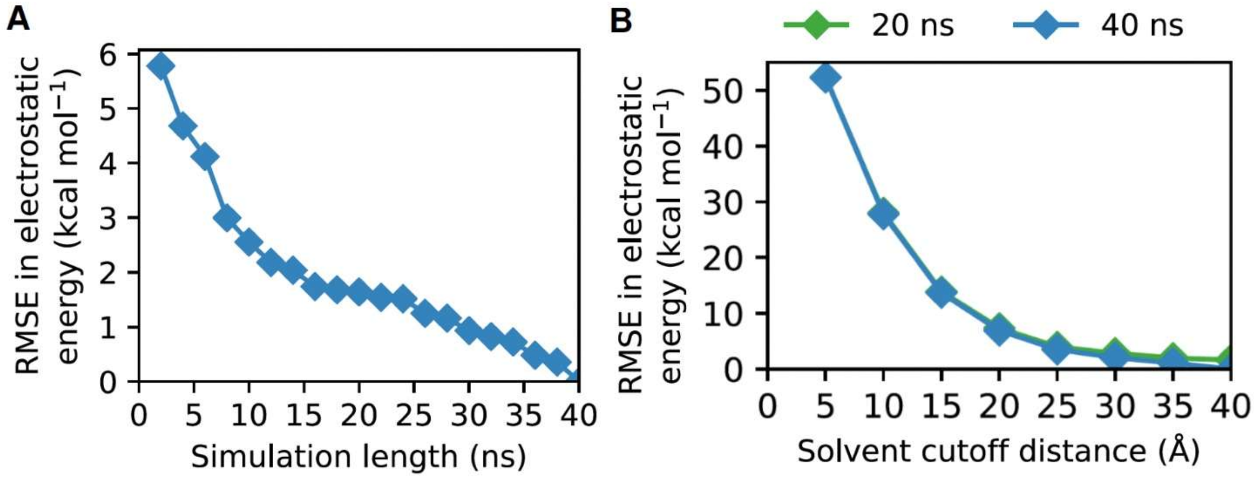


Figure S4: Convergence of electrostatics as a function of simulation length and box size. (A) RMSE in electrostatic energy (ESE) across all the RNA atoms in the validation set conformations for different simulation lengths. The solvent cutoff was 40 Å. (B) RMSE in the ESE across all the validation set conformations with different solvent cutoff distances. The time-averaged ESE is computed with 20 ns (green) and 40 ns (blue) simulation lengths.

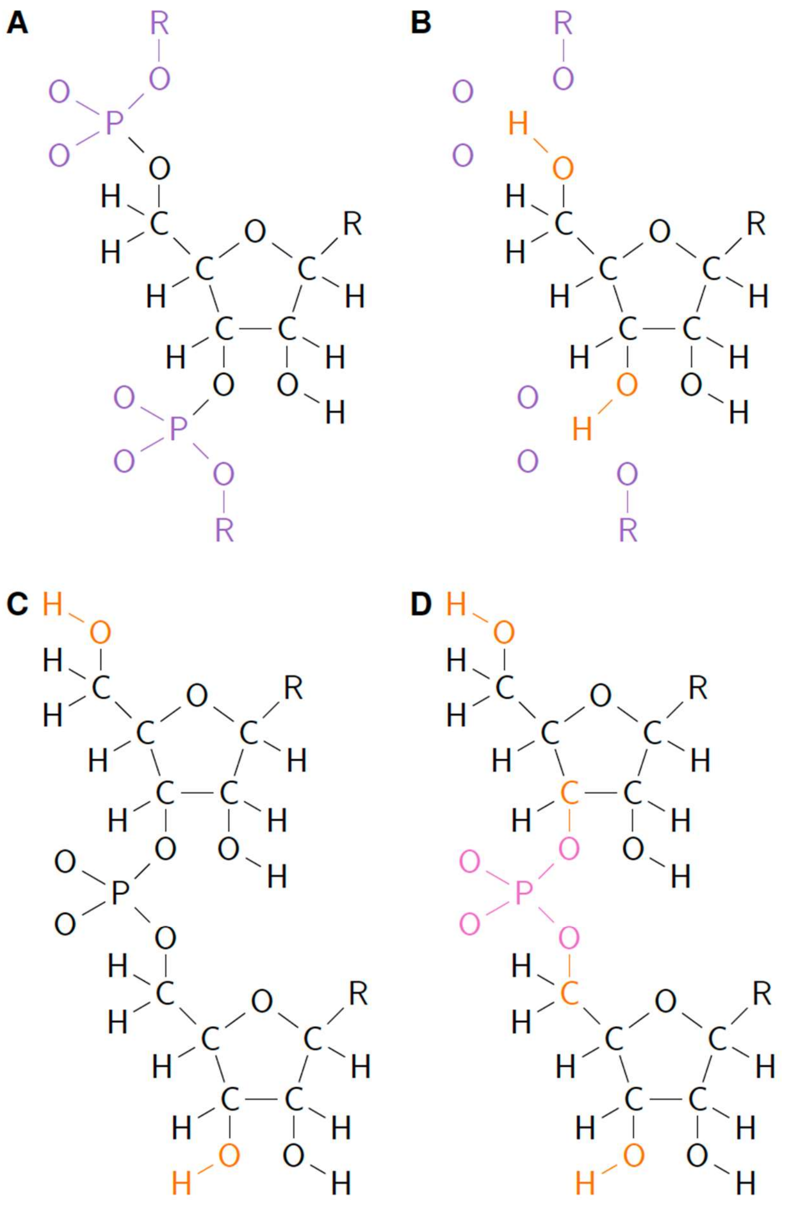


Figure S5: QM atom embedding scheme for nucleotides. (A) For non-adjacent residues, the phosphate atoms were treated as MM atoms; QM atoms are in black; MM atoms are in purple. (B) Closed shell representation for non-adjacent residues used in SAPT calculations. The MM atoms were represented as point charges that are not covalently bonded to an interacting RNA residue. The OH covalent bond was oriented in the same direction as OP and is marked in orange. (C) The embedding scheme for adjacent interacting residues (or for base-phosphate interaction). (D) The closed shell representation for adjacent interacting residues. The linking phosphate is marked in pink.

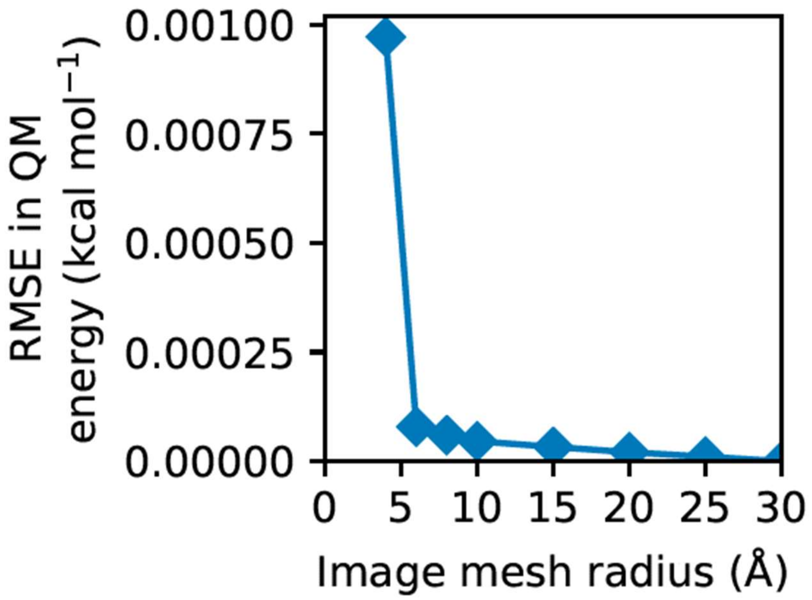


Figure S6: RMSE of QM interaction energy across different conformations in the validation set as a function of image mesh radii. All solvent atoms beyond the image mesh were replaced by image charges on the mesh. The reference was a QM calculation with point charges representing all solvent atoms within 30 Å of any RNA atom.

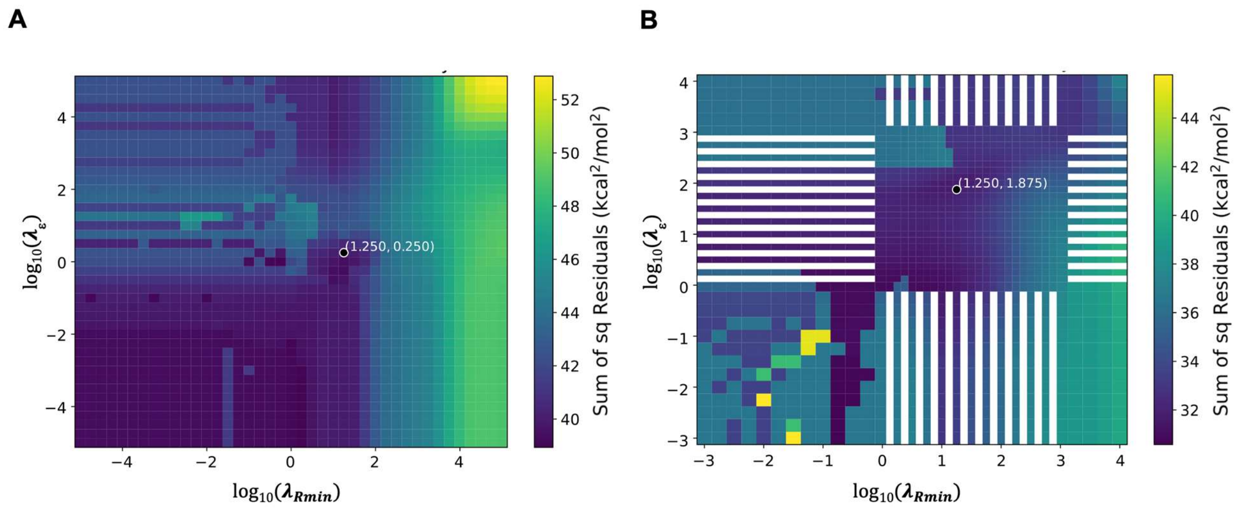


Figure S7: Minimum cross validation errors for iteration 1 (Panel A) and iteration 2 (Panel B). This shows a larger area that was tested than the regions shown in Figure 5. For iteration 2, some grid values were not tested and are shown as white in the plot. The model is prone to overfitting at lower $\lambda$ values. The chosen optimal regularization strength is marked on each heatmap. The error difference between the selected regularization strength and minimum error regularization strength is ~1-2 $kcal^2/mol^2$.

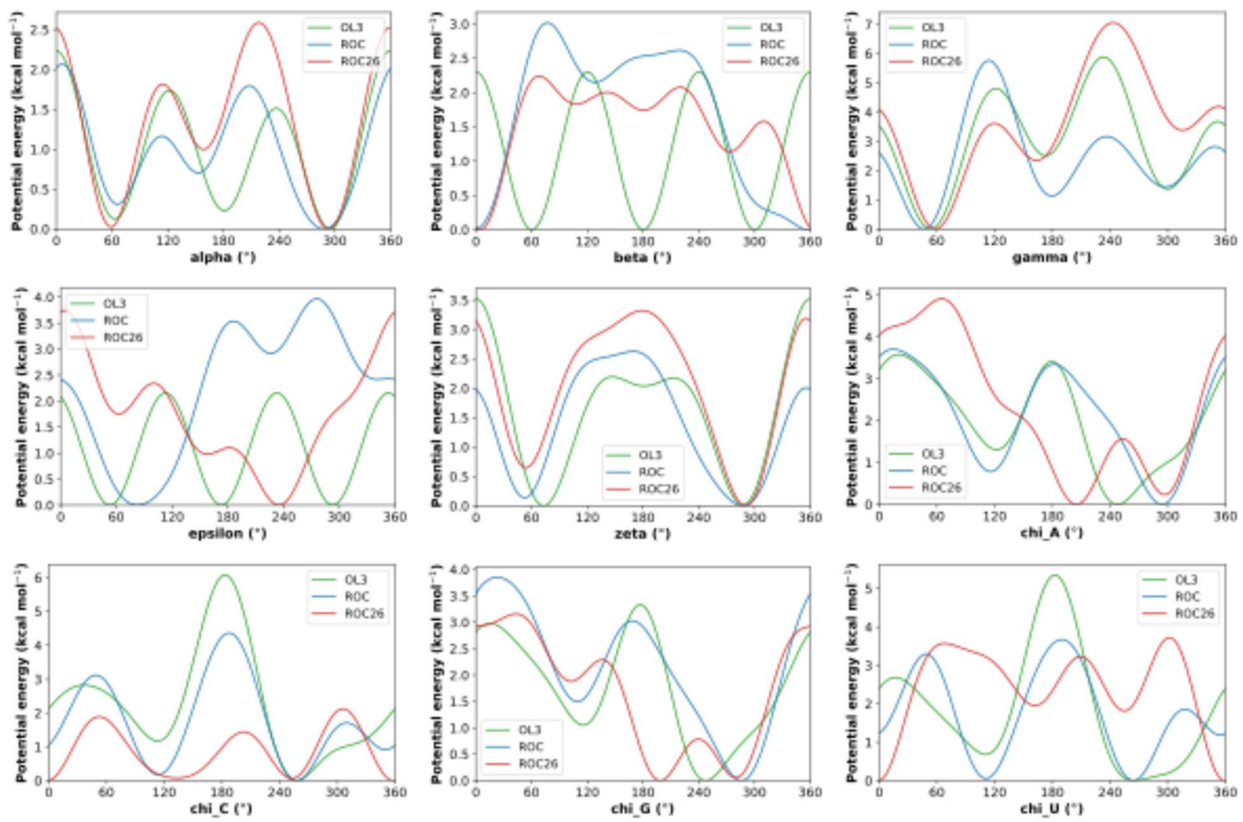


Figure S8: Dihedral potential energy profile using new dihedral parameters. The new dihedral parameters are obtained using the RNA.ROC26 nonbonded parameters with the torsion fitting method of Aytenfisu et al[1]. The profiles are compared to the current Amber force field (OL3)[2] and to RNA.ROC[1].

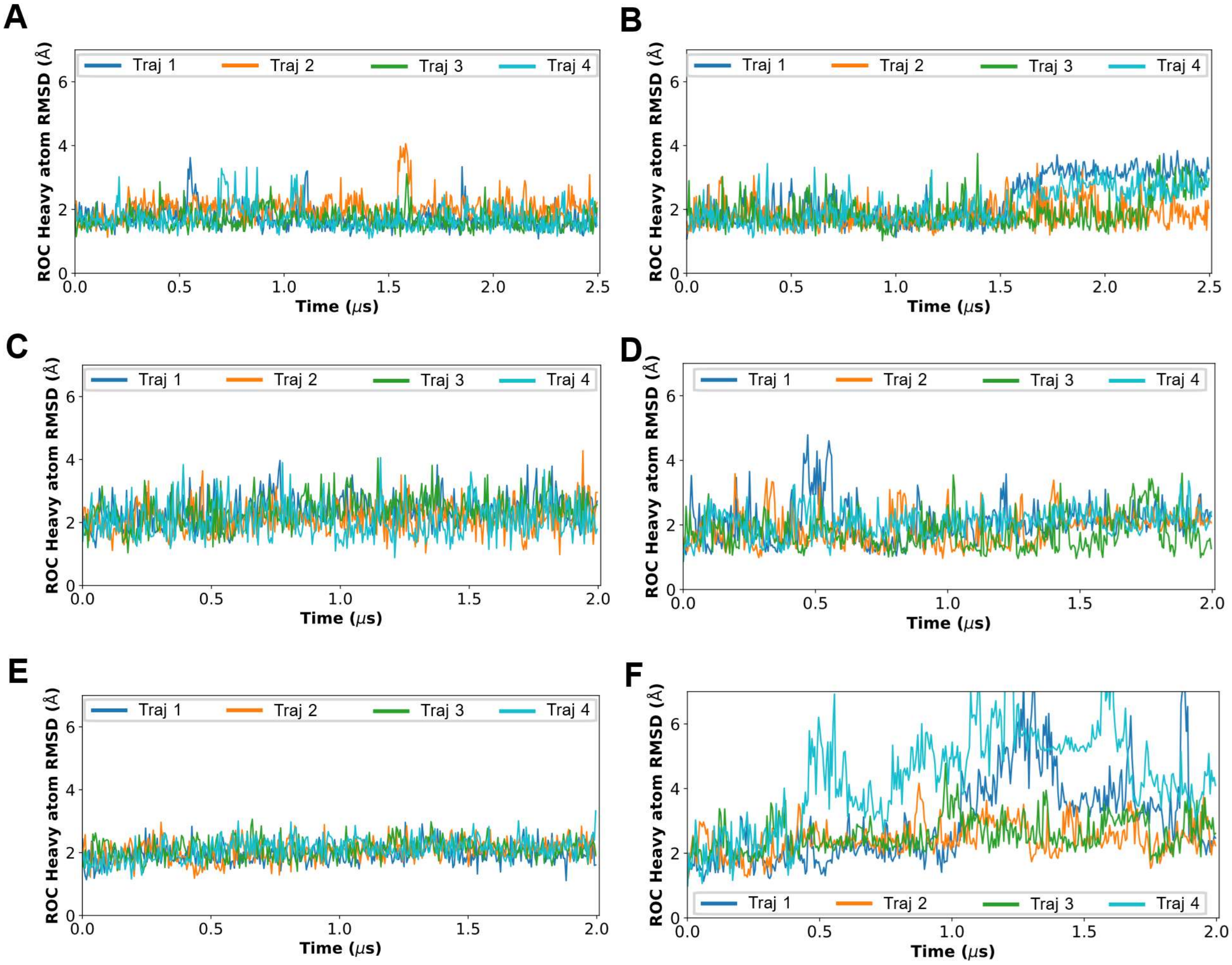


Figure S9: RMSD as a function of time for 2 μs simulation of 1ZIJ (Panel A), 2KOC (Panel B), 2JXQ (Panel C), GC Duplex (Panel D), 2DD2 (Panel E) and AU Duplex (Panel F) using the RNA.ROC[1] force field.

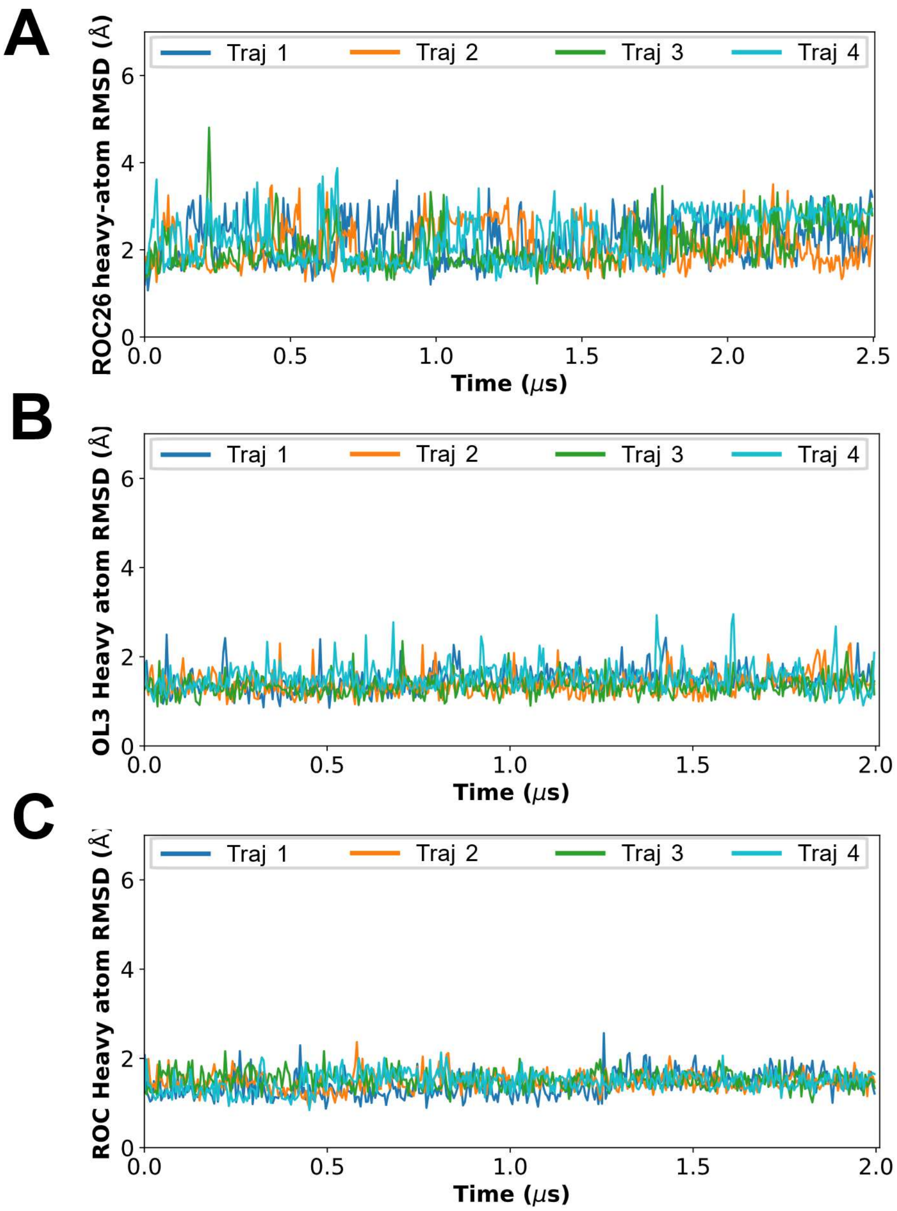


Figure S10: RMSD as a function of time for the internal loop including closing base pairs for 2DD2 (5'GGUGAAGGCC/3' CCGAAGCCG 5'). Panel A is for RNA.ROC 26 forcefield (2.5 µs), Panel B for Amber ff99 + bsc0 + $\chi_{OL3}$ force field (2 µs) and Panel C for RNA.ROC (2 µs). The larger fluctuations in RNA.ROC26 are a result of A5 moving away from the internal loop.

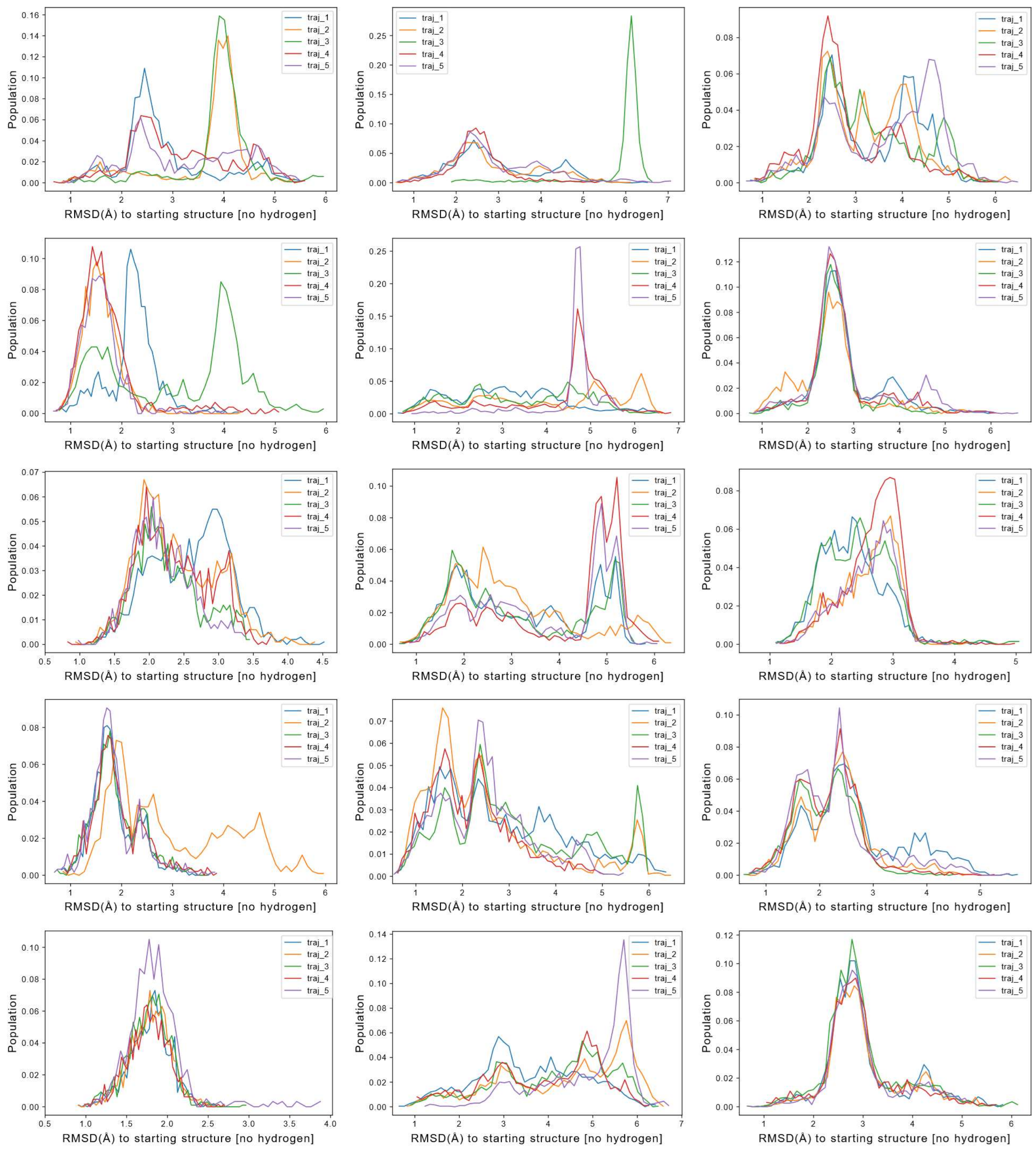


Figure S11: Histogram of RMSD to the A-form reference structure made using NAB. From top to bottom are sequences: AAAA, CAAU, CCCC, GACC, and UUUU. Results from RNA.ROC26 MD simulations are on the left, RNA.OL3 (the current Amber force field) in the middle and RNA.ROC on the right. Five independent simulations were run for each tetramer.

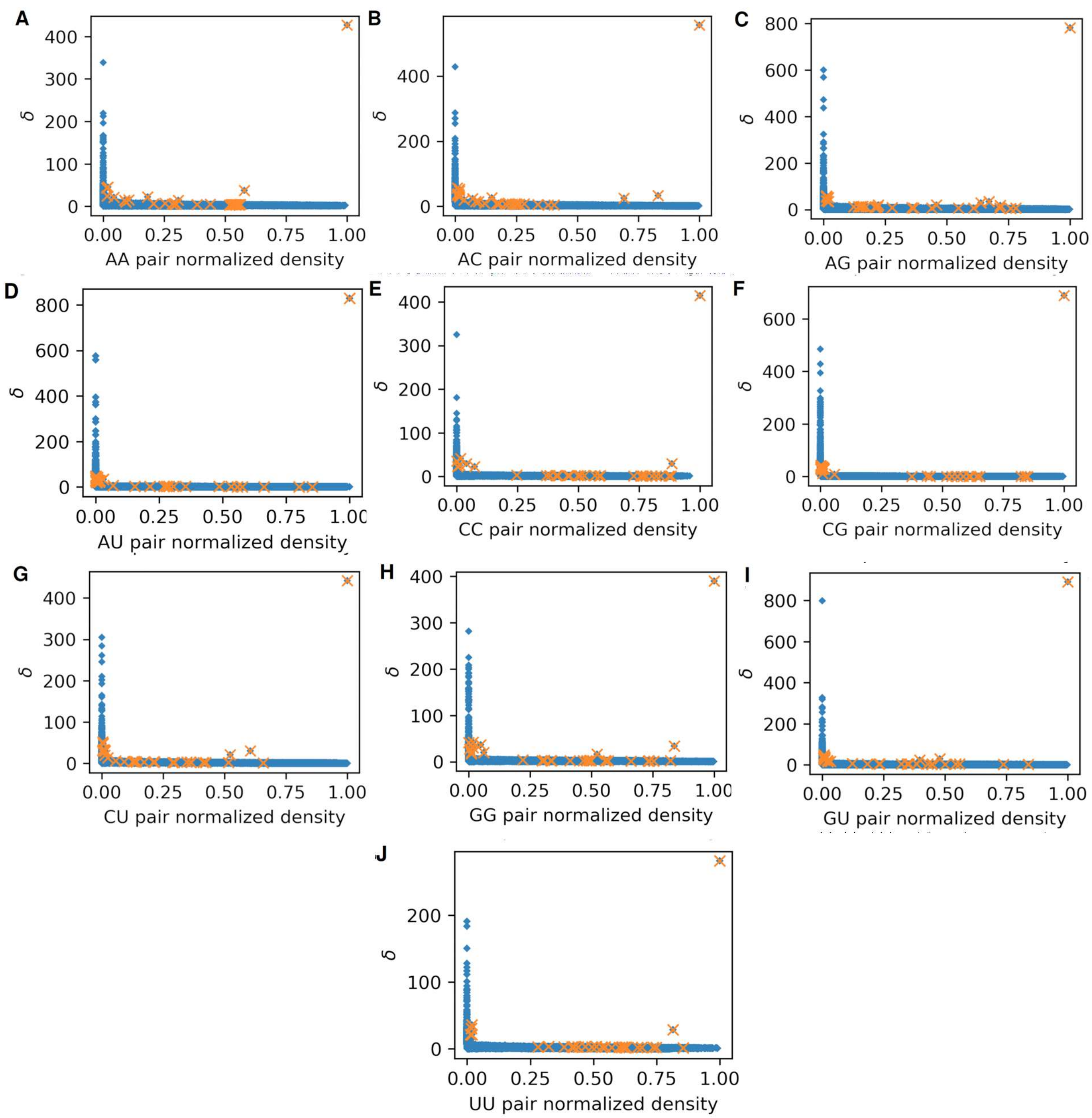


Figure S12: Decision plots from density peak clustering of pairs of interacting RNA residues by nucleobase identity. The minimum distance from any point with a higher density $\delta$ is plotted against normalized density. The selected 32 structures are marked in orange for each RNA pair type. All other candidate conformations are marked in blue.
